# The Rpd3 histone deacetylase is critical for temperature-mediated morphogenesis and virulence in the human fungal pathogen *Histoplasma*

**DOI:** 10.1101/2025.08.01.668069

**Authors:** Nebat Ali, Mark Voorhies, Rosa A. Rodriguez, Anita Sil

**Affiliations:** Department of Microbiology and Immunology, University of California, San Francisco, San Francisco, California, USA; Chan Zuckerberg Biohub – San Francisco, San Francisco, California, USA

## Abstract

Adaptive responses to environmental stimuli are integral to the survival and virulence of microbial pathogens. The thermally dimorphic human fungal pathogen *Histoplasma* senses temperature to transition between a mold form in soil and a pathogenic yeast in mammalian hosts. The contributions of chromatin-modifying enzymes to the ability of *Histoplasma* to appropriately respond to temperature have never been explored. Through chemical inhibition and genetics, we determined that the class I histone deacetylase (HDAC) *RPD3* is required for normal *Histoplasma* yeast morphology at 37 °C. Rpd3 regulated the expression of key morphology-specific genes, including critical virulence factors and transcription factors (TFs), was required for normal DNA-binding activity of yeast-promoting TFs, and influenced histone acetylation levels at the loci of putative pro-filamentation TFs. Furthermore, Rpd3 was required for virulence in a macrophage model of infection. Taken together, Rpd3 is a critical regulatory component that both activates the pathogenesis program and represses the filamentation program to enable thermal dimorphism in *Histoplasma*. This work uncovers the crucial role that chromatin regulation plays in temperature response of this ubiquitous pathogen.

## Introduction

The ability to sense and effectively adapt to mammalian hosts is a hallmark of clinically relevant microbial pathogens. This phenomenon can be observed in thermally dimorphic fungi, such as the primary human pathogen *Histoplasma*, where sensing of elevated host body temperature triggers a dramatic shift in cell state that enables growth and persistence within the host. *Histoplasma* is a globally distributed pathogen and has been identified on all seven continents [1–3]. In the United States, it is endemic to the Ohio and Mississippi River Valleys where it is estimated to infect nearly 250,000 individuals annually [4]. In the soil environment at ambient temperature (or in the laboratory at room temperature (RT)), *Histoplasma* grows as a filamentous mold and produces infectious spores that can aerosolize. Spores and hyphal fragments can be inhaled into the lungs of a mammalian host, where elevated body temperature (37 °C) is sufficient to trigger formation of the parasitic budding yeast form. *Histoplasma* yeast produce a wide arsenal of secreted effectors to bypass host innate immune cells in the lungs and cause the acute pulmonary disease histoplasmosis [5–10, 65–67, 102–105]. In severe cases, acute histoplasmosis may develop into a chronic pulmonary disease that can result in a life-threatening disseminated infection [1].

The regulatory mechanisms driving the ability of *Histoplasma* to sense and respond to temperature have been a key area of study. Prior transcriptomics studies to characterize temperature-regulated morphology in *Histoplasma* found that up to a quarter of the transcriptome changes in response to growth at 37 °C vs RT [11–13]. Additional studies showed that the yeast program at 37 °C is dependent on the activity of interdependent TFs that are <u>R</u>equired for <u>Y</u>east <u>P</u>hase growth (*RYP1-4*) [15–17]. The Ryp TFs accumulate at 37 °C and cooperate to direct the expression of yeast-phase genes [15–17]. Mutants lacking Ryp TFs grow constitutively as hyphae at 37 °C and display transcriptional signatures that are virtually indistinguishable from wild-type cells grown at RT [17]. Studies aimed at further characterizing the regulatory mechanisms that direct morphogenesis in *Histoplasma* also identified the TFs *WET1, STU1, FBC1,* and *PAC2* for their roles in promoting filamentation [18–19, 106].

While there have been key advances in identifying TFs that regulate *Histoplasma* thermal dimorphism, additional regulatory mechanisms remain largely unexplored. The capacity of TFs to exert their regulatory ability can be dictated by how accessible the genome is for TF binding and the recruitment of transcriptional machinery [21–22] but it is unknown if regulation of chromatin state by histone-modifying enzymes influences temperature-dependent gene expression and the activity of temperature-responsive TFs in *Histoplasma.* Studies from fungi to vertebrates have identified roles for histone modifying enzymes such as histone deacetylases (HDACs) and histone acetyltransferases (HATs) in coordinating genome accessibility to facilitate large-scale changes in gene expression and TF activity in response to environmental changes [23–26]. There are numerous examples of this type of regulation in model fungi [27–30], human fungal pathogens [31–34], and fungal pathogens of plants [35–38].

In this study, we explored HDAC-mediated mechanisms that could contribute to gene regulation in *Histoplasma.* We focused our study on classical zinc-dependent HDACs due to their widespread functions in virulence, metabolism and morphogenesis in a diverse set of fungal pathogens [26, 38–39]. By employing a combination of small-molecule HDAC inhibitors and targeted genetics, we uncovered a novel role for the Rpd3 HDAC in *Histoplasma* morphology and gene expression in response to host temperature. Chemical inhibition and genetic ablation of *RPD3* at 37 °C globally disrupted the genome-wide binding patterns of the temperature-responsive Ryp TFs. Furthermore, loss of *RPD3* altered histone acetylation patterns of candidate pro-filamentation genes, corresponding with their transcriptional activation. Lastly, we found that Rpd3 is required for virulence in a macrophage model of infection. Taken together, we identified and characterized Rpd3 as a key chromatin regulator that integrates with the temperature-responsive Ryp network to regulate a critical developmental switch in *Histoplasma*.

## Materials and Methods

### Ethics statement

All animal work was approved under UCSF Institutional Animal Care and Use Committee protocol # AN181753-02B

### *Histoplasma* strains and culture conditions

All experiments and strain manipulations were conducted with *Histoplasma ohiense* G217B unless explicitly indicated. Strains used in this study are listed in Table S1. For studies involving G217B and different geographical *Histoplasma* isolates, we used G217B (ATCC 26032), G184AR (ATCC 26027) and G186AR (ATCC 26029) strains generously provided by the laboratory of William Goldman at the University of North Carolina, Chapel Hill and sourced HcH88 (ATCC 32281) from the American Type Culture Collection (ATCC). For TF mutant studies, we used strains generated in Joehnk *et al.* 2023 [53] and Assa *et al.* 2023 [19] derived from the parental *G217Bura5^-^* WU15 strain that was also provided by the Goldman laboratory. Growth of liquid *Histoplasma* cultures under host conditions was carried out at 37 °C and 5% CO_2_ with shaking in Histoplasma Macrophage Media (HMM) supplemented with Penicillin/Streptomycin (P/S). RT growth conditions were carried out at ambient temperature between 22-24 °C with shaking in HMM containing an equimolar amount of GlcNAc (A3286, Sigma Aldrich) instead of glucose as previously described (Gilmore *et al.* 2013). All HDACi treatment experiments conducted at 37 °C were conducted in HMM GlcNAc media. For HDACi growth assays, SAHA (LC laboratories V-8477), Entinostat (MedChem Express HY-12163), Mocetinostat (Selleck Chemicals S1122), Ricolinostat (TargetMol T2489), MC1568 (Cayman Chemical Company 16265), and TMP159 (Selleck Chemicals S8502) were ordered as 10 mM solutions in DMSO or as powders that were resuspended to a concentration of 10 mM in DMSO (Sigma-Aldrich D8418) before being filter sterilized using a 0.22 µM MCE filter (Celltreat 229751) and diluted further in sterile DMSO for growth assays.

### Culture conditions for HDACi assays

Experiments where cultures are supplemented with an additive (DMSO or HDACi) followed the general scheme outlined in Figure 1B and were conducted as described previously [19]. Cultures used for growth assays, RNA-seq and ChIP-seq studies were set up in triplicate biological replicates. When possible, inoculum from a single colony grown at 37 °C was used to start a liquid culture in HMM media. This liquid culture was passaged every 2-3 days in HMM media for a total of 2 passages before being used to inoculate a master culture in HMM + GlcNAc at an OD_600_ of 0.2 that was then aliquoted into replicates with equal volumes of each HDACi concentration being tested or DMSO. Cultures were incubated for 2 days at 37 °C and 5 % CO_2_ or at RT with shaking. Microscopy samples were harvested and fixed for 30 minutes with a solution of 4 % paraformaldehyde (PFA) every 24 hours for up to 3 days following incubation and stored long-term at 4 °C in PBS. Samples for genomic DNA isolation were harvested through centrifugation. Cell pellets used for isolating RNA, and protein samples were harvested through filtration using SFCA Membrane Nalgene Rapid-Flow Sterile Disposable Bottle Top Filters (Fisher 09-740-22G). 10-25 mL of culture was harvested from each flask and immediately flash frozen in liquid nitrogen before being stored at -80 °C.

**Figure 1.**
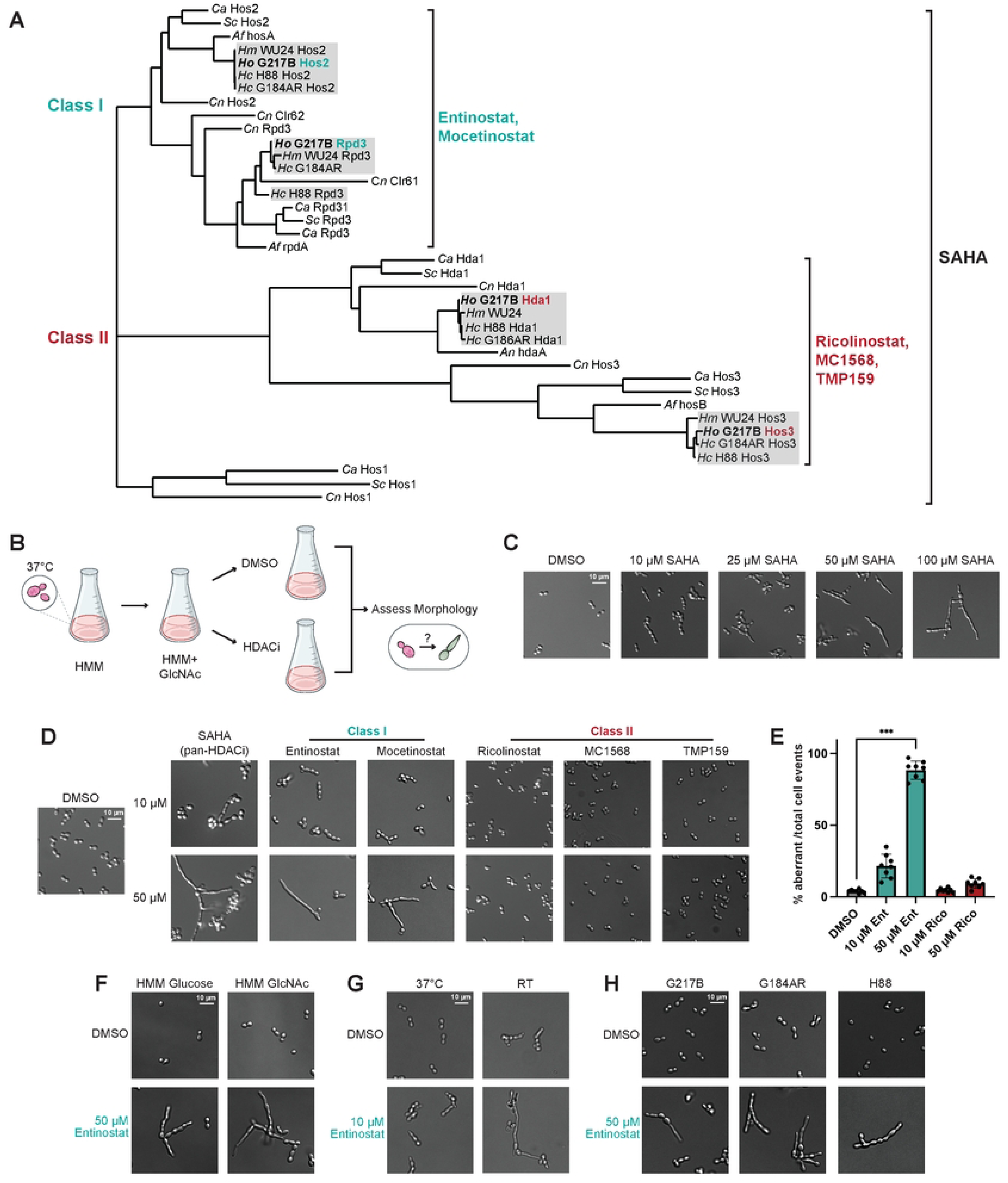
Disrupting conserved HDACs in *Histoplasma* promotes improper filamentation at 37 °C. A. Phylogenetic tree of classic HDAC domain-containing proteins in *Histoplasma spp.* (G217B, G186AR, H88, and WU24) and a panel of representative fungi that includes *S. cerevisiae, C. albicans, A. fumigatus,* and *C. neoformans*. Colors distinguish the two main HDAC classes. HDACs found in the G217B strain primarily used throughout this study are bolded. Single brackets indicate the targeting range of each set of chemical inhibitors. B. Schematic outlining the assay used to assess *Histoplasma* morphology following exposure to chemical HDAC inhibitors. Micrographs displaying *Histoplasma* G217B morphology following growth at 37 °C for 2 days in (C) the presence of DMSO or an increasing concentration of the pan-HDAC inhibitor SAHA or (D) DMSO or 10 µM and 50 µM of a class-specific HDAC inhibitor. E. Quantification of abnormal (non-yeast) cell morphology events in DMSO, Entinostat, and Ricolinostat-treated samples represented in (D). Each datapoint represents n >100 cell events. F. Micrographs displaying *Histoplasma* G217B morphology following growth at 37 °C for 2 days in the presence of 50 µM of Entinostat in media supplemented with glucose or GlcNAc. G. Images of cells treated with 10 µM Entinostat for 2 days following passage at 37 °C or RT. H. Assessment of cell morphology in different geographical species of *Histoplasma* grown for 2 days at 37 °C with 50 µM of Entinostat. *** p < 0.001, Wilcoxon rank-sum test.

### Cloning and mutant generation

Mutant and OE strains in this study were generated in *Histoplasma ohiense* G217B (ATCC 26032). Plasmids used in this study are listed in Table S2. Constructs were generated in *E. coli* DH5⍺ (New England Biolabs (NEB) C2988J) and OneShot TOP10 (Fisher Scientific C404003) cells. CRISPR/Cas9 plasmids for targeted KO were generated as previously described [53]. Protospacer sequences flanking the *RPD3* and *HOS2* loci were selected using a local version of CHOPCHOP [81]. Partial sgRNA cassettes were ordered as gBlocks (Table S4) from Integrated DNA Technologies (IDT) and pairs of 5’ and 3’ targeting sgRNA cassettes were assembled into pNA16, a pDONR/Zeo entry clone containing an sgRNA insertion site in between the *Histoplasma* glyceraldehyde-3-phosphate dehydrogenase (GAPDH) promoter and *A. nidulans* tryptophan synthase terminator (trpC) using Gibson Assembly (NEB E2611S). We used Gateway LR clonase (Thermo Fisher Scientific 11791100) to introduce the fully sgRNA expression cassette assembled in the donor plasmid into pBJ262, a hygromycin (*hph)* marked Cas9 episomal expression plasmid used for transformation in *Histoplasma.* Final plasmids were sequence verified for correct assembly and linearized using *PacI* (NEB R0547L) along with a non-targeting control plasmid before being electroporated into yeast as described previously [53].

Strains were screened on solid HMM media supplemented with 200 µg/mL of Hygromycin B (Life Technologies, Inc./Gibco Laboratories 10687-010) to select for positive transformants. Transformants were initially screened via colony PCR (cPCR) using primers (see Table S3) that fall outside of the sgRNA cut sites. Promising KO isolates were passaged on solid media with selection and subjected to additional cPCR screening using primers internal to one of the sgRNA cut sites. In some cases, single cell resuspensions of transformants were plated for additional cPCR screening. Transformants displaying a clean KO via flanking or internal cPCR were then grown in liquid HMM and passaged 3-4 times in the absence of hygromycin to facilitate plasmid loss. Following plasmid loss, these strains were then subjected to genomic DNA extraction and whole genome sequencing to validate successful KO (outlined below).

*RPD3* overexpression plasmids were generated through amplifying the *RPD3* open reading frame (ORF) from genomic DNA using Q5 Polymerase (NEB M0494S) and primers (see Table S3) overlapping with a pDONR overexpression backbone amplified from pDI71. The primers used mediate insertion of the *RPD3* ORF in between the *ACT1* promoter and *CATB* terminator sequences derived from pDI71 via NEB HiFi Assembly cloning (NEB E5520S). To generate an OE plasmid containing the putative catalytic dead variant of *RPD3,* site-directed mutagenesis (SDM) on the WT entry plasmid (pNA61) was carried out using the Agilent Quickchange Lightning SDM Kit (Agilent Technologies 210518). SDM primers used were informed by the residue mapping outlined in Figure S4A. The resulting *RPD3* OE pDONR plasmids were cloned into the *hph*-marked pSB238 episomal plasmid and validated via whole plasmid sequencing prior to transformation and selection on HMM + hyg solid media. Whole plasmid sequencing for cloning was performed by Plasmidsaurus using Oxford Nanopore Technology with custom analysis and annotation.

### *Histoplasma* Imaging

PFA-fixed samples were diluted in PBS and mounted on a glass-bottom 96-well imaging plate with high performance #1.5 cover glass (Cellvis P96-1.5H-N). Micrograph images were taken using a 40X DIC objective lens on a Zeiss Axiovert 200m microscope.

### RNA Extraction

Total RNA was extracted from fungal cells as previously described [18–19] with Qiazol (Qiagen 79306). Frozen cell pellets were resuspended in equal volumes of Qiazol and subjugated to bead beating with 0.5 µM zirconia beads (Spectrum Chemical 140-19990-E1) and extraction using chloroform. The resulting aqueous phase was transferred to Epoch SpinTM RNA columns (Epoch Life Sciences 1940-250) to isolate RNA and subjected to sequential washes with 3 M NaOAc (pH = 5.5) and 10 mM TrisCl (pH = 7.5) in 80 % ethanol. The columns were then treated with Purelink DNAse (Invitrogen 12185010) and subjected to additional washes with 3 M NaOAc (pH = 5.5) and 10 mM TrisCl (pH = 7.5) in 80 % ethanol before being eluted with RNAse-free water. The organic phases of each sample were reserved at -20 °C for total protein extraction.

### mRNA Isolation

Equal amounts of total RNA (between 5-10 ng) across all samples were used for mRNA isolation. These samples were subjected to a second round of DNAse treatment (Turbo DNAse, Fisher Scientific AM1907) and their quality was assessed with an RNA 6000 Nano Bioanalyzer Kit (Agilent Technologies 5067-1511). Three rounds of poly-A selection using oligo-dT Dynabeads® (Invitrogen 61005) was used to purify mRNA, and mRNA enrichment was verified with an RNA 6000 Nano Bioanalyzer chip.

### RNA-seq library preparation

poly-A-selected RNA was used to generate libraries with the NEB Next Ultra II Directional RNA Library Prep Kit for Illumina (NEB E7760S). Each sample was dual indexed with unique primers (UCSF Center For Advanced Technology (CAT)) compatible for multiplexed sequencing. Library quality and size was assessed with the Agilent High Sensitivity DNA Bioanalyzer Kit (Agilent Technologies 5067-4626) to ensure proper fragment size (∼300-400 bp) and the absence of contaminating adapters. Final library concentrations were measured using a Qubit HS dsDNA assay kit (Life Technologies Q33230) and equal amounts were pooled into a final library for sequencing. Pooled libraries were used for paired-end sequencing using one NextSeq 2000 lane at the Chan-Zuckerberg Biohub (Fig. 2) or single-end sequencing using one NovaSeq X lane at the UCSF CAT (Fig. 4).

**Figure 2.**
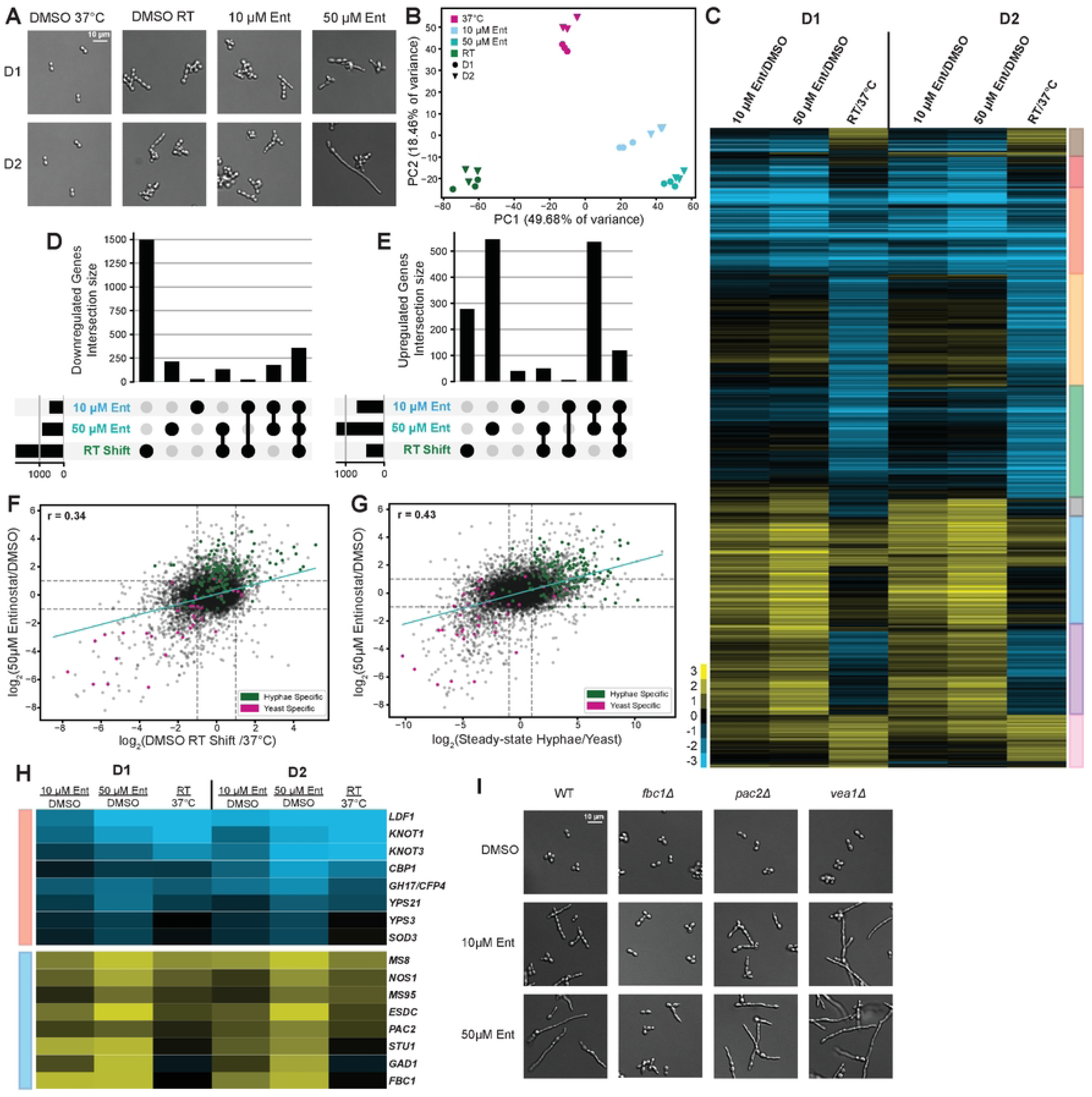
Temperature-mediated transcription at 37 °C relies on class I HDAC activity. A. Representative images of cells used in transcriptomics studies. Samples were obtained from the same flask, 24 hours apart across all replicates. B. Principal component analysis (PCA) plot generated using counts per million (CPM) values of all transcripts across the triplicate samples represented in A. C. Heatmap displaying 4241 differentially expressed genes passing significance (LIMMA-fit > 2-fold change, 5% FDR) in cells grown with Entinostat at 37 °C or at RT at early timepoints. Rows represent unique transcripts, and enrichment is calculated as the log_2_ CPM ratio of each Entinostat or RT sample to the time-matched yeast control at 37 °C. log_2_ CPM ratio values are indicated by heatmap coloring. The vertical color bar to the right of the heatmap indicates gene groups defined through bi-clustering. UpSet plots highlighting the shared and unique differential gene sets 2 days following RT shift and Entinostat-treatment that are (D) negatively and (E) positively regulated. Scatterplots comparing differential RNA-seq signal in all G217B transcripts between day 2 RT and 50 µM Entinostat samples (F) and day 2 50 µM Entinostat samples and steady-state hyphae (G) (Gilmore *et al.* 2015). Pearson’s correlation coefficient is denoted by *r*. H. Zoomed-in view of heatmap clusters identified in C to highlight differential yeast-phase specific (YPS, top panel) and hyphal-phase specific (HPS, bottom panel) genes of interest. I. Micrographs of *fbc1Δ*, *pac2Δ*, and *vea1Δ* strains [19, 53] following 2 days of treatment with 10 µM and 50 µM of Entinostat. “WT” in this instance refers to the parental WU15 (*G217Bura5^-^)* strain used for generating *fbc1Δ*, *pac2Δ*, and *vea1Δ* [19, 53].

**Figure 3.**
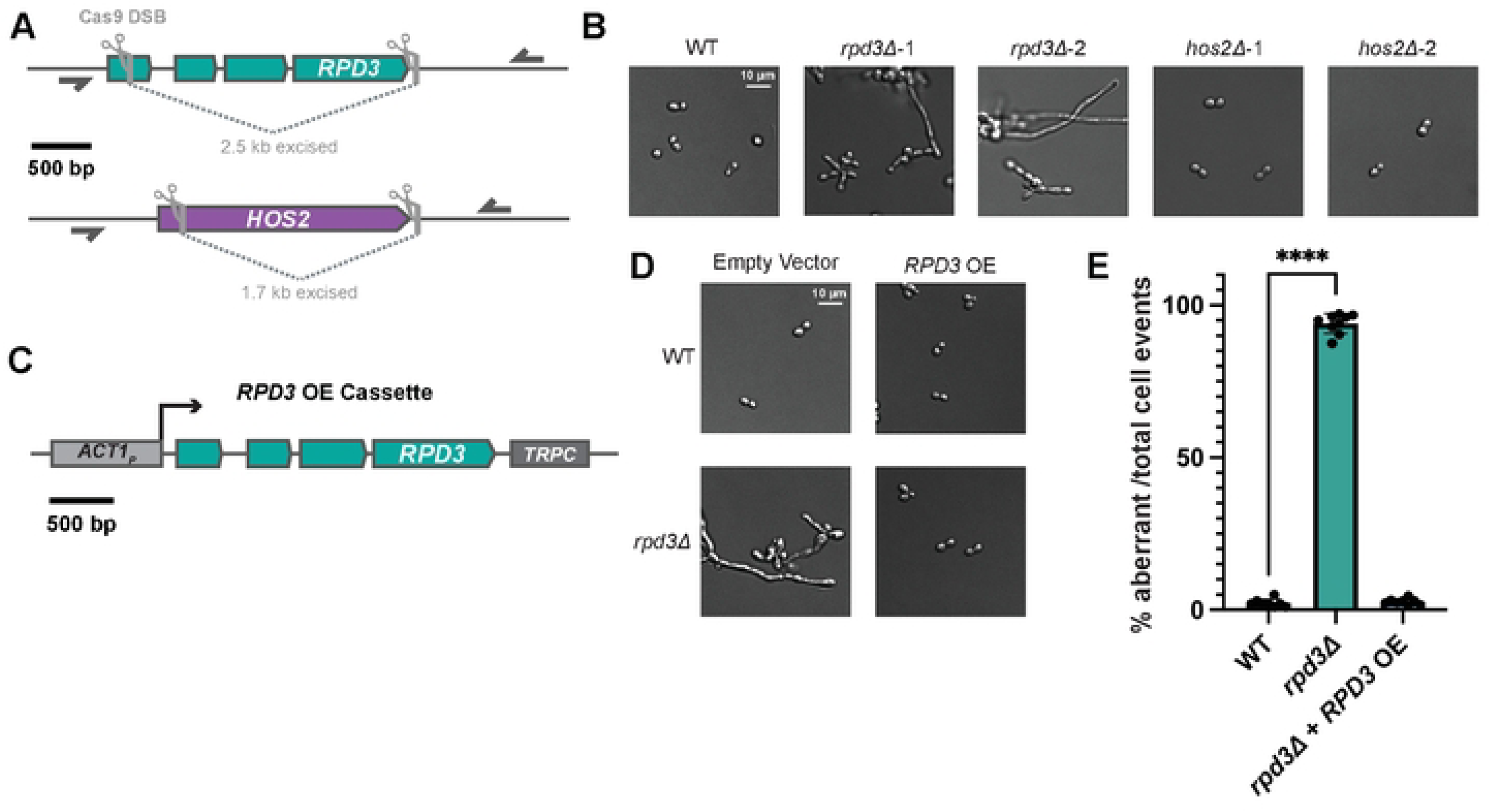
Using CRISPR/Cas9 to generate class I HDAC mutants reveals *RPD3* is required for yeast growth at 37 °C. A. Schematic illustrating *RPD3* and *HOS2* loci that were targeted for KO using sgRNA-guided Cas9. sgRNA sites that undergo double-stranded breaks (DSBs) by Cas9 are indicated by vertical gray lines. The horizontal half-arrows indicate PCR primers used to assess removal of the region between the two sgRNA sites. B. Representative cell morphology images of G217B control (WT) cells alongside two *rpd3Δ* and *hos2Δ* isolates grown at 37 °C. C. Schematic illustrating overexpression cassette used to drive expression of wild-type *RPD3* under the *ACT1* promoter. D. Micrographs of WT G217B and *rpd3Δ* strains transformed with an empty vector control or a construct overexpressing WT *RPD3* at 37 °C. E. Quantification of abnormal (non-yeast) cell morphology events for WT, *rpd3Δ*, and *rpd3Δ* + *RPD3* OE samples represented in (B) and (D). Each datapoint represents n >100 cell events. *** p < 0.0001, Wilcoxon rank-sum test.

**Figure 4.**
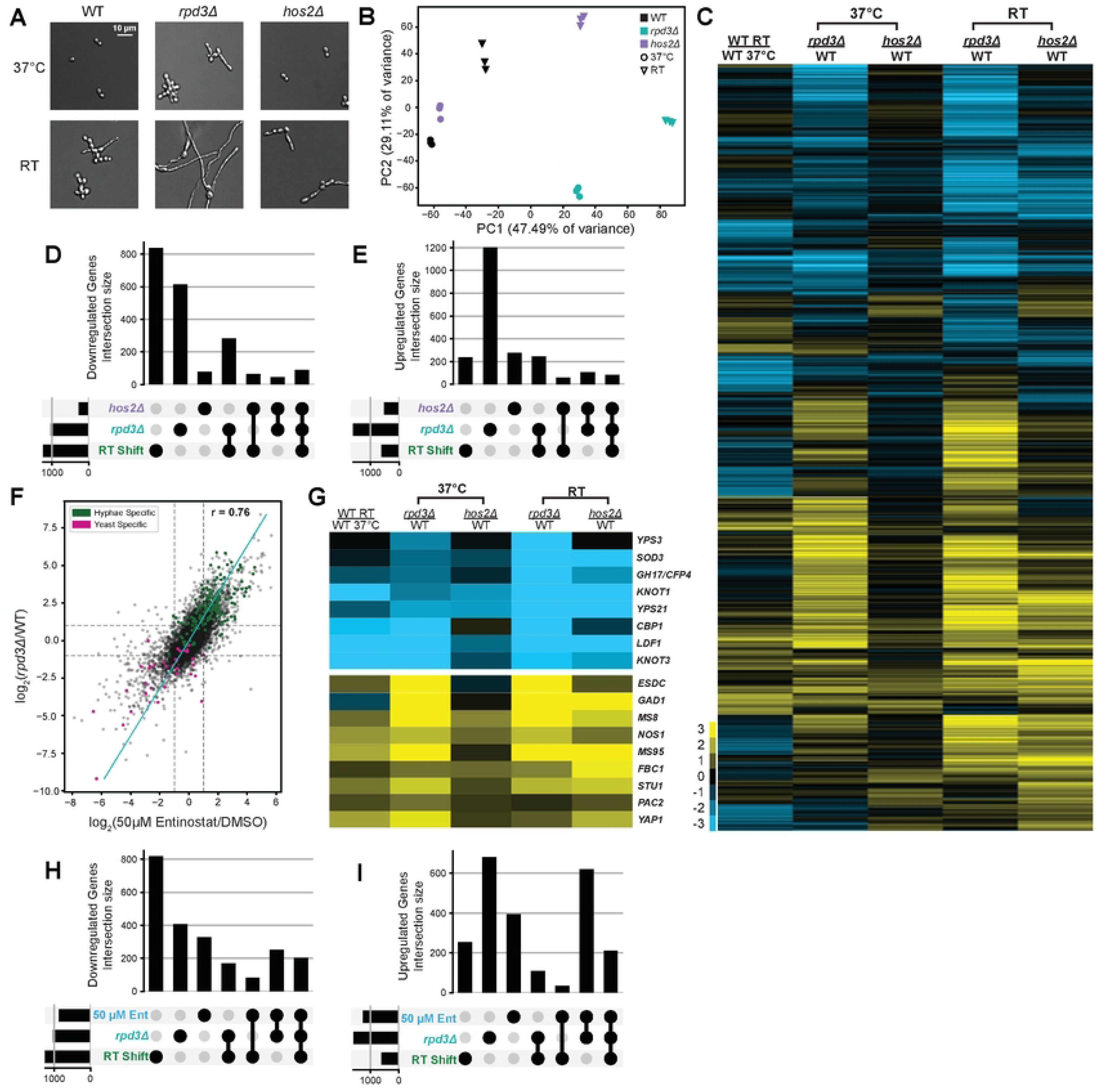
Transcriptomic analysis of *rpd3Δ* reveals global disruption to phase-specific gene expression. A. Representative micrographs of cells used for transcriptomics studies. G217B control (WT), *rpd3Δ,* and *hos2Δ* were passaged and grown in parallel at 37 °C in HMM and at RT in HMM + GlcNAc for 2 days. B. Principal component analysis (PCA) plot generated using counts per million (CPM) values of all transcripts expressed across the samples represented in (A). C. Heatmap of 5480 differentially expressed genes passing significance (LIMMA-fit > 2-fold change, 5 % FDR) in *rpd3Δ* and *hos2Δ* strains at 37 °C and RT relative to WT controls at each respective temperature. Differentially expressed genes in WT samples at 37 °C and RT were included in this analysis for comparison. Ratios are displayed as log_2_ CPM values that are indicated by heatmap color. UpSet plots displaying the intersections of differentially expressed genes identified in *rpd3Δ* and *hos2Δ* strains at 37 °C or in WT cells following 2-day RT shift that are transcriptionally repressed (D) or upregulated (E). F. Scatterplot comparing global differential RNA-seq signal in *rpd3Δ* to the signal in cells following 2-day treatment with 50 µM Entinostat. Pearson’s correlation coefficient is denoted by *r*. G. Expanded view of heatmap in (C) to highlight Yeast-Phase Specific (YPS-top panel) and Hyphal-Phase Specific (HPS, bottom panel) genes of interest that are significantly dysregulated in *rpd3Δ*. UpSet plots listing the differentially expressed gene sets found in *rpd3Δ,* 2-day RT shift and 2-day 50 µM Entinostat treatment to identify downregulated (H) and upregulated (I) genes that are shared and unique across strains and conditions.

### RNA-seq analysis

Transcript abundances were quantified based on version ucsf_hc.01_1.G217B of the *Histoplasma* G217B transcriptome (S5 Data of Gilmore *et al.* 2015 [18]). Relative abundances (reported as TPM values [82] and estimated counts (est_counts) of each transcript in each sample were estimated by alignment free comparison of k-mers between the reads and mRNA sequences using KALLISTO version 0.46.2 [83]. Further analysis was restricted to transcripts with estimated counts ≥ 10 in at least four samples (for the experiment in Figure 2) or at least nine samples (for the experiment in Figure 4). Differentially expressed genes were identified by comparing replicate means for contrasts of interest using LIMMA version 3.46.0 [84–85]. Genes were considered significantly differentially expressed if they were statistically significant (at 5% FDR) with an effect size of at least 2x (absolute log_2_ fold change ≥ 1) for a given contrast.

### Protein extraction

Organic phases reserved at -20 °C were washed with 100% ethanol and spun down to pellet DNA and isolate the protein-containing phenol-ethanol supernatant. Proteins were precipitated from this suspension using isopropanol and subjected to washes with 0.3 M guanidinium thiocyanate in 95 % ethanol before being resuspended in 100 % ethanol. Protein pellets were then spun down and dried at RT before being dissolved in freshly-made urea lysis buffer (9 M Urea, 25 mM Tris-HCl, 1% SDS, 0.7 M β-mercaptoethanol) and boiled for 5 minutes. Protein concentrations were measured using a Pierce 660 nm Assay Kit (Thermo Scientific 22660) supplemented with the ionic detergent compatibility reagent (Fisher Scientific PI22663).

### Western Blotting

Total protein extracts were obtained from pellets used to conduct the ChIP-seq and RNA-seq studies outlined in Figs. 4-5. 10 µg of total protein resuspended in Novex 554 NuPAGE LDS Sample Buffer (Invitrogen NP0007) was boiled for 5 minutes, loaded on NuPAGE Novex 4-12% Bis-Tris Gels (Life Technologies Corporation NP0322BOX) and subjected to electrophoresis with MOPS buffer at 140 V. Proteins were subjected to semi-dry transfer on nitrocellulose membranes using the Invitrogen iBlot2 Gel transfer device. Membranes were incubated with Intercept® (PBS) Blocking Buffer (LI-COR) for one hour before being incubated in custom primary polyclonal antibodies raised against Ryp1 (ID: 3873), Ryp2 (ID: 387), and Ryp3 (ID: 356) [15–17] or a primary antibody against tubulin (Novus Biologicals DM1A) at 4 °C with shaking overnight. Following incubation, membranes were washed three times with a solution of 0.1 % Tween-20 in PBS (PBST) before being incubated with LICOR Infrared Dye-conjugated secondary antibodies (IRDye 800CW goat ⍺-Rabbit IgG for rabbit ⍺-Ryp antibodies and Donkey ⍺-Mouse IgG for mouse ⍺-tubulin antibody) for one hour at RT. Membranes were subjected to three washes in 0.1 % PBST before being visualized using the LI-COR Odyssey CLx imaging system.

**Figure 5.**
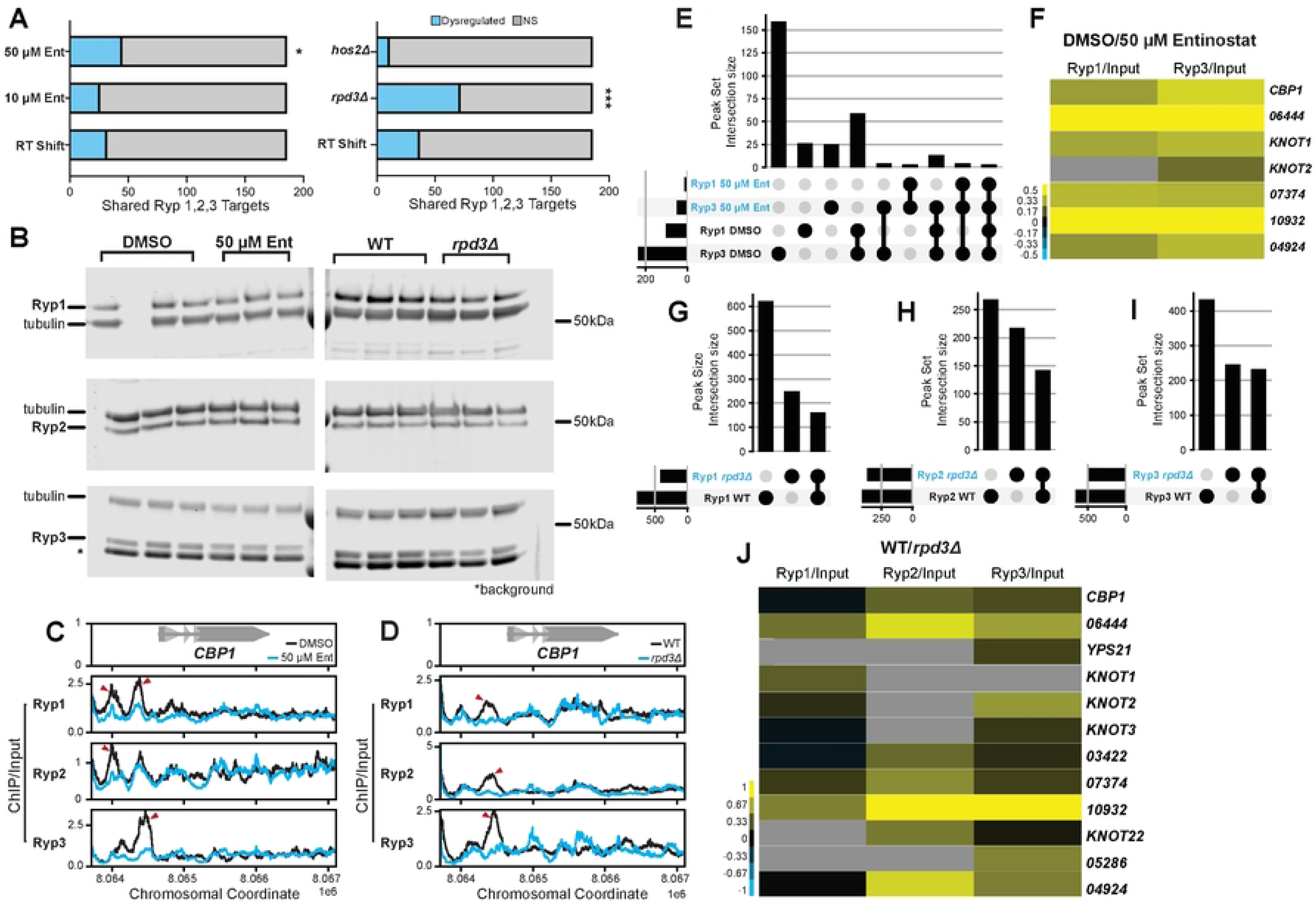
The activity of temperature-responsive TFs is dependent on Rpd3. A. Stacked box plots listing the number of shared Ryp1-3 regulatory targets [17] that are dysregulated following 2-day growth at RT, Entinostat treatment, and KO of class I HDAC genes. Statistical significance calculated by Fisher’s Exact is indicated by asterisks: *=p<0.05, ***=p<0.001. B. Western Blots performed on total protein samples extracted from WT G217B cells grown for 2 days at 37 °C with DMSO or 50 µM Entinostat and WT G217B or *rpd3Δ* strains grown for 2 days at 37 °C. Whole protein extracts were derived from the cell pellets used for the transcriptomics studies outlined in Figs. 2 and 4, respectively. The asterisk indicates background signal observed with the Ryp3 antibody. Coverage tracks displaying fold enrichment traces of ChIP/input signal for each Ryp pulldown at the locus surrounding *CBP1* in WT G217B cells following 2 days of growth at 37 °C with DMSO or 50µM Entinostat (C) and G217B WT or *rpd3Δ* strains grown for 2 days at 37 °C (D). E. UpSet plot displaying the intersecting sets of genes with Ryp-binding events in their promoter regions following DMSO or 50 µM Entinostat treatment for 2 days at 37 °C. F. Heatmap displaying enrichment of Ryp1 and Ryp3 occupancy in peaks found upstream of putative virulence genes (defined in Rodriguez *et al.* 2025 [20]) in WT cells grown with DMSO or 50 µM Entinostat for 2 days at 37 °C. Heatmap colors display the fold enrichment of ChIP/input signal between the two conditions. UpSet plots displaying genes in WT and *rpd3Δ* cells with promoter binding events for Ryp1 (G), Ryp2 (H), and Ryp3 (I). J. Heatmap displaying quantification of Ryp promoter occupancy in the promoters of virulence genes as shown in (F) for ChIP signal observed in WT and *rpd3Δ* strains grown for 2 days at 37 °C.

### Detecting secreted proteins in *Histoplasma*

Secreted proteins were detected as previously described [9]. G217B WT and *rpd3Δ* cells cultured under standard host conditions were grown to saturation before equal volumes were pelleted through centrifugation. The resulting supernatants were filter sterilized using a 0.22 µM syringe filter and subjected to concentration using Amicon Ultra Centrifugal Filter Units with size cutoff of 3 kDa cutoff (Millipore Sigma UFC900324). For each sample, equal amounts of concentrated supernatants were separated using SDS-PAGE and stained using InstantBlue Coomassie Protein Stain (ISB1L –abcam 119211). Gels were then washed with PBS and imaged using a Canon Canoscan LiDE210.

### Genomic DNA isolation and sequencing in *Histoplasma*

Cell pellets for genomic DNA (gDNA) isolation were harvested through pelleting 1 mL of culture via centrifugation at 12000 rpm. Cells were washed and resuspended in 1X TE before being subjected to DNA extraction using the Qiagen Gentra Puregene Yeast/Bacteria kit (158567) as previously described [20]. Purified gDNA was treated with RNase A before being quantified and submitted for library preparation using the Illumina DNA prep kit and Whole Genome Sequencing (WGS) at SeqCenter. Paired-end (PE) sequencing was performed using an Illumina NovaSeq X Plus at SeqCenter.

### Whole genome sequencing analysis

Reads were aligned to the *Histoplasma ohiense* G217B genome (GCA_017607445.1) [86] using BWA MEM [87]. We confirmed deletion of target genes and lack large CNV by visual inspection of read coverage.

### Chromatin isolation and shearing

Cultures processed for ChIP-seq were crosslinked with formaldehyde (Polysciences 18814-10) added to a final concentration of 1 % and kept under culture conditions (shaking at 37 °C with 5 % CO_2_ or ambient RT) for 20 minutes to allow fixing to take place. Crosslinking was quenched with glycine added to a final concentration of 125 mM for 5 minutes at RT. Fixed cells were pelleted at 3000 g for 10 minutes at 4 °C and washed twice with TBS (20 mM Tris-HCl (pH 7.5), 150 mM NaCl). Washed cells were harvested through filtration, aliquoted into pellets of equal mass (between 150-300 mg/tube) and flash frozen in liquid nitrogen as described above.

Frozen pellets were resuspended in lysis buffer (50 mM Hepes/KOH (pH 7.5), 140 mM NaCl, 1 mM EDTA, 1 % Triton X-100, 0.1 % sodium deoxycholate, protease/phosphatase inhibitors) at 4 °C before being bead beat with 500 µL of 0.5 mm zirconia beads in 5 x 1 minute cycles at RT with 2 minute rests on ice in between. The resulting lysate was then spun down at 8000 rpm for 10 minutes at 4 °C to isolate the insoluble chromatin fraction. This chromatin pellet was resuspended in 600-900 µL of lysis buffer and sonicated at 4 °C using the Diagenode Bioruptor® Pico for 30-35 cycles (30s on, 30s off). Sonicated samples were spun down for 5 min at 14,000 rpm at 4 °C to pellet cell debris and aliquots for input DNA were reserved at -20 °C in TE + 1 % SDS. Input samples were also used to assess shearing using a High Sensitivity DNA Bioanalyzer chip from Agilent. Equal lysate volumes were aliquoted for each IP to be performed and brought to a final volume of 500 µL with lysis buffer and 20 µg of each polyclonal ⍺-Ryp antibody (Ryp1: 3873, Ryp2: 387, and Ryp3: 356) or 3 µg of a ChIP-grade monoclonal ⍺-histone-acetyl antibody (H3K9Ac: ab177177, H3K14Ac: ab52946, H4K16Ac: ab109463). Samples were incubated overnight with agitation at 4 °C before being incubated with 50 µL of a 50% slurry of Protein A Dynabeads® (Life Technologies Corporation 10002D) pre-washed 2x with TBS and 3x lysis buffer for 2 hours at 4 °C with agitation.

Immunoprecipitated DNA was recovered through pulldown on Protein A beads using a magnetic rack. Pellets containing DNA-protein complexes were washed 2x in lysis buffer, 2x with high-salt (500 mM NaCl) lysis buffer, 2x with wash buffer (10 mM Tris/HCl (pH 8.0), 250 mM LiCl, 0.5 % NP-40, 0.5 % Na-deoxycholate, 1 mM EDTA) and 1x with TE before adding elution buffer (50 mM Tris/HCl (pH 8.0), 10 mM EDTA, 1 % SDS) and incubating at 65 °C for 10 minutes. DNA was eluted using a magnetic rack and the resulting beads were subjected to second round of resuspension and elution with 150 µl TE + 0.65 % SDS and pooled with the initial eluate to maximize total yield of DNA. IP and input samples were treated with 1 µL of Proteinase K (20 mg/mL Qiagen 19131) and incubated at 65 °C for approximately 16 hours. IP and Input DNA samples were then treated with 2 µg of RNase A (Qiagen) and incubated at 37 °C for 1 hour. DNA was then purified using the Zymo ChIP DNA Clean & Concentrator Kit (D5205) and eluted in 31 µL of ddH_2_O or TE.

### ChIP-seq library preparation

Purified IP and input DNA were used for Illumina library preparation using the NEBNext Ultra II DNA Library Prep Kit (NEB E7103S) and Illumina dual-indexed primer pairs sourced from the UCSF CAT. The quality and size distribution of each library was assessed using the Agilent High Sensitivity DNA Bioanalyzer Kit (Agilent Technologies 5067-4626) to ensure proper fragment size and the absence of contaminating adapters. Final library concentrations were measured using a Qubit HS dsDNA assay kit (Life Technologies Q33230) and used to pool equimolar amounts of each sample into final libraries for sequencing. Pooled libraries were used for single-end sequencing (DMSO vs 50 µM Ent samples in Figures 5 and 6) or paired-end sequencing (WT vs *rpd3Δ* samples in Figures 5 and 6) using one NovaSeq X 10B lane at the UCSF CAT.

**Figure 6.**
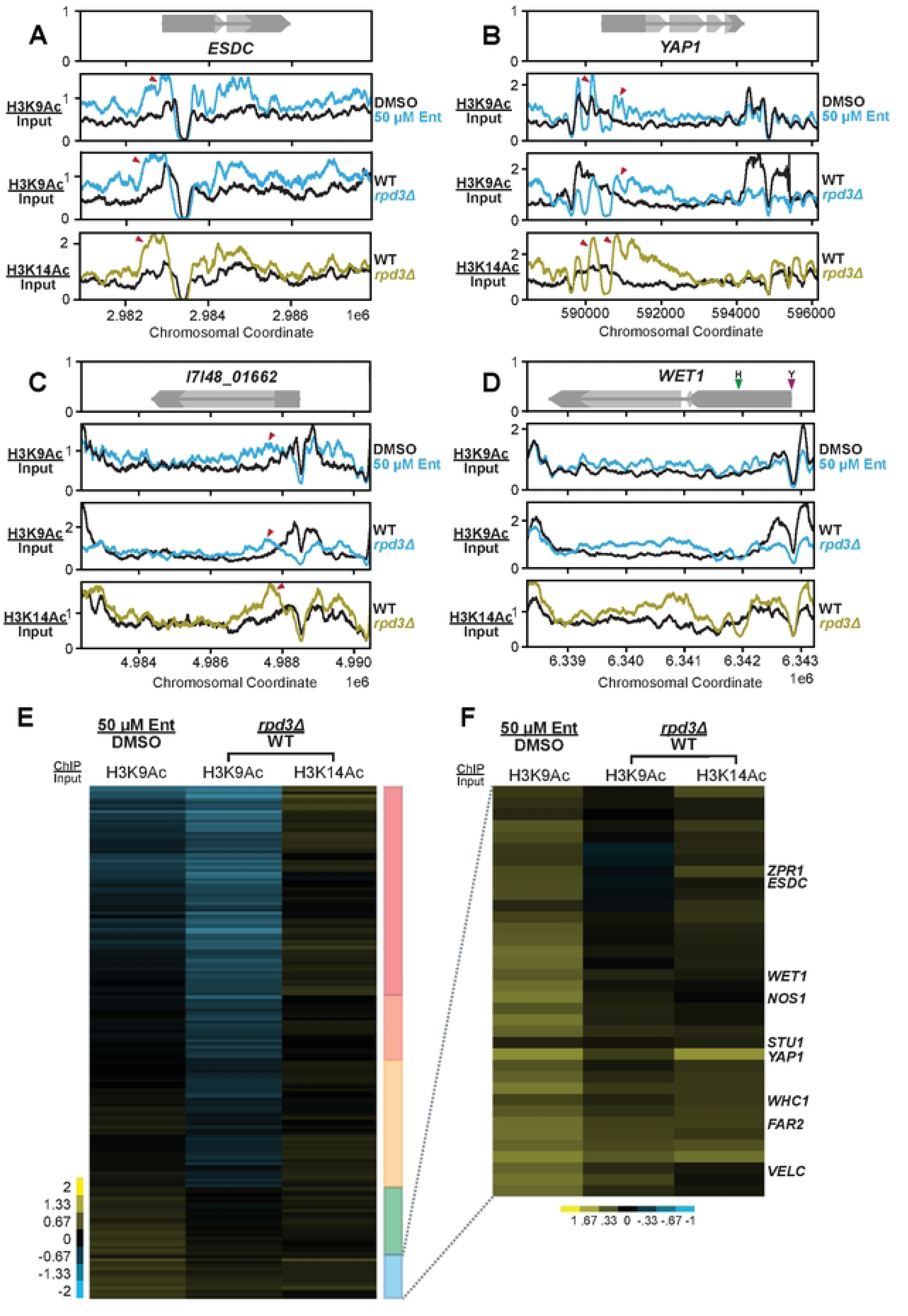
Rpd3 drives histone H3 hypoacetylation at the loci of hyphal TF genes at 37 °C. Coverage tracks displaying traces of genomic ChIP/input signal for pulldowns on two different histone H3 acetylation marks, H3K9Ac and H3K14Ac. WT yeast treated with DMSO or 50 µM Entinostat along with WT and *rpd3Δ* strains grown for 2-days at 37 °C were used for chromatin isolation and subsequent pulldown. Colored traces (H3K9Ac in red and H3K14Ac in green) display signal in control (DMSO, WT) cells while the black traces represent signal with Entinostat or in *rpd3Δ* surrounding the loci of *ESDC* (A), *YAP1* (B) and *I1I48_01662* (C). The gene prediction schematic in the top panel shows coding sequences (CDS) in light gray and untranslated regions (UTRs) in dark grey. D. Same as in (A-C) for the locus surrounding *WET1.* Colored arrows on the gene prediction schematic highlight the start sites of two different UTR isoforms enriched for this transcript in yeast (Y, purple) and hyphae (H, green). E. Sorted heatmap displaying fold enrichment of histone-acetyl ChIP signal for H3K9Ac and H3K14Ac in the promoters of putative transcription factors encoded across the G217B genome. F. Expanded heatmap view to highlight differential H3K9Ac and H3K14Ac signal within a cluster of hyphae-associated TFs. Heatmap colors indicate differential ChIP/input signal intensity across the two comparisons.

### Transcription factor ChIP-seq TF analysis

Single or paired-end reads were aligned to the *Histoplasma ohiense* G217B genome (GCA_017607445.1) [86] using BOWTIE2 version 2.4.2 [88] with default parameters. Peaks were called for each individual replicate with MACS2 version 2.2.7.1 [56] with default parameters except that min-length was set to 100 for paired-end reads as recommended by Gaspar 2018. Aggregate peaks were defined by taking the union of the coordinates of overlapping peaks among replicates for regions where peaks were called for at least two replicates [90]. The aggregate peaks defined for 37 °C samples for a given TF were used for quantification of all samples for that TF. Per peak signal was quantified by counting reads or read pairs in each aggregate peak, log_2_ transforming with pseudocount of 1 to handle missing data, and normalizing to the signal in the *NPS1* negative control region. This negative control region was defined as the middle 500 bp of *NPS1* (*I7I48_05472,* JAEVHH010000002: 444765..449764 (+)). Differential enrichment was then calculated as the difference of the means of the log_2_ signal over replicates from each condition. Peaks were assigned to individual genes if any part of the peak fell in the intergenic region between the stop codon of the upstream gene and before the start codon for that gene, limiting the intergenic size to a maximum of 10 kb as described in [90].

### Histone modification ChIP-seq analysis

Single or paired-end reads were aligned to the *Histoplasma ohiense* G217B genome (GCA_017607445.1) [86] using BOWTIE2 version 2.4.2 [88] with default parameters. Per sample promoter signal was quantified by counting reads or read pairs in each promoter, log_2_ transforming with pseudocount of 1 to handle missing data, and normalizing to the signal in the *NPS1* negative control region. This negative control region was defined as the middle 500 bp of *NPS1* (*I7I48_05472,* JAEVHH010000002: 444765..449764 (+)). Differential enrichment was then calculated as the difference of the means of the log_2_ signal over replicates from each condition.

### Phylogenetic analysis and tree generation

HMMSEARCH from HMMER 3.3.2 [91] was used to query the Hist_deacetyl (PF00850) Pfam domain against predicted gene sets for 41 fungal genomes (S18 Data of Gilmore *et al.* 2015 [18]) yielding 172 proteins with expectation value ≤ 1e-6. These sequences were aligned with PROBCONS 1.12 [92] and a phylogenetic tree was inferred from the alignment using FASTTREE 2.1.11 [93].

### Other Software and Libraries

We wrote custom scripts and generated plots in R 4.0.4 [94] and Python 3.9.2, using Numpy 1.19.5 [95] and Matplotlib [96]. Jupyter 4.7.1 [97] notebooks and JavaTreeView 1.1.6r4 [97] were used for interactive data exploration.

### BMDM differentiation and culture conditions

Bone marrow derived macrophages (BMDMs) were isolated from the tibias and femurs of 6-8 week-old C57BL/6J (Jackson Laboratories stock, No. 000664) mice. Mice were euthanized via CO_2_ narcosis and cervical dislocation as approved under UCSF Institutional Animal Care and Use Committee protocol. Cells were differentiated in bone marrow macrophage media (BMM) containing Dulbecco’s Modified Eagle Medium (DMEM High Glucose), 20 % Fetal Bovine Serum (FBS), 10 % v/v CMG supernatant (the source of CSF-1), 2 mM glutamine, 110 mg/mL sodium pyruvate, 110 mg/mL penicillin and streptomycin. Undifferentiated monocytes were plated in BMM for 7 days at 37 °C with 5 % CO_2_. Adherent cells were collected and frozen in 40% FBS and 10 % DMSO and stored in liquid nitrogen for future use.

### Macrophage infections

Macrophage infections with G217B Histoplasma strains were performed as previously described [7–9, 102]. Macrophages were seeded in tissue culture-treated dishes in triplicate one day before infection and incubated at 37 °C with 5% CO2. On the day of infection, yeast cells from logarithmic-phase cultures (OD_600_ = 4-7) were collected, resuspended in D-PBS (PBS Ca^2+^, Mg^2+^ free), sonicated for 3 seconds on setting 2 using a Fisher Scientific Sonic Dismembrator Model 100, and counted at 40X using a hemacytometer. Due to the morphology phenotype of the *rpd3Δ* mutant, chains of budding yeast cells were counted as one cell. Macrophages were infected at appropriate MOIs and allowed to phagocytose *Histoplasma* for 2-hours before being washed with D-PBS to remove any extracellular yeast and fresh BMM media was added. For infections lasting longer than 2 days, the remaining cells were supplemented with fresh BMM media.

### Lactate dehydrogenase (LDH) release assay

To quantify macrophage lysis, BMDMs were plated (7.5 x 10^4^ cells per well of a 48-well plate) and infected in triplicate as described above. At the indicated time points, the amount of LDH in the supernatant was measured as described previously [99–100] BMDM lysis was calculated as the percentage of total LDH from supernatant of wells with uninfected macrophages lysed in 1 % Triton X-100 at the time of infection. Due to continued replication of BMDMs over the course of the experiment, the total LDH at later time points can be greater than the total LDH from the initial time point, resulting in lysis that is greater than 100 %.

### Immunofluorescence of *rpd3Δ* infection

To image macrophages, BMDMs were seeded at 1.5 x 10^5^ cells on 12 mm circular glass coverslips in 24-well tissue culture dishes. Infections were done as previously described with G217B wild type and the *rpd3Δ* mutant *Histoplasma*. At each time point post infection, cells were fixed with 1 mL of 4 % paraformaldehyde for 20 minutes at room temperature. After fixation, coverslips were washed 3 times with D-PBS to remove any residual fixative. Fixed cells were permeabilized and blocked using 0.2 % w/v Saponin and 2 % FBS in D-PBS for 1 hour at room temperature. Cells were then incubated with AlexaFluor-647-conjugated anti-F4/80 mouse antibody (UCSF mAB core) at a dilution of 1:500 for 1 hour at 37 °C in a light-blocking incubation chamber. Coverslips were washed 5 times with D-PBS and then briefly rinsed in double distilled water. Excess liquid was gently removed using edge of a Kimwipe. Finally, coverslips were mounted on glass slides with ∼10 µL of Vectashield mounting media with DAPI and 1:100 Calcofluor White (1 mg/mL). Coverslips were sealed to the glass slide using clear nail polish.

### Image acquisition and processing

Z-stacks (0.3 mm increments) images were taken on Nikon Eclipse Ti CSU-X1 spinning disk confocal microscope. FIJI [101] was used to generate max-intensity projections for all conditions.

## Results

### Chemical inhibition of histone deacetylases disrupts *Histoplasma* thermal dimorphism

To identify HDAC-encoding genes in *Histoplasma*, we used a Hidden Markov Model of the classical HDAC domain (PF00850, PMID 9278492) to query a panel of fungal proteomes that included four representative species of *Histoplasma.* The phylogenetic tree generated by this analysis revealed two distinct classes of Zn^2+^-dependent HDACs in *Histoplasma* (Fig. 1A). We used reference sequences from *Saccharomyces cerevisiae* to assign *Histoplasma* orthology corresponding to Class I (Rpd3 and Hos2) and Class II (Hos3 and Hda1) HDACs. Expanding this analysis to include a broader set of environmental and pathogenic fungal species highlighted the shared conservation and minimal expansion of these two HDAC classes across fungi (Fig. S1A). While paralogs of the Class I HDAC *RPD3* are found in opportunistic fungal pathogens such as *C. albicans* and *C. neoformans* [31, 34], we did not identify any expansion of HDACs in the panel of thermally dimorphic fungal pathogens included in this analysis (Fig. S1A). The Hos1 outgroup found in *S. cerevisiae*, *Candida spp.* and *C. neoformans* [40] was similarly absent in *Histoplasma spp.* (Fig.1A) in addition to the other fungal genomes included in this analysis (Fig. S1A).

To investigate a putative role for histone deacetylases (HDACs) in *Histoplasma* thermal dimorphism, we assessed the morphology of yeast cells following challenge with small-molecule inhibitors of classical (i.e Zn^2+^ dependent) HDACs. Because functional redundancy of classic HDACs has been observed in related fungal species [34, 41–42], we initially tested the effect of Suberoylanilide Hydroxamic Acid (SAHA), a pan-HDAC inhibitor capable of targeting all classic HDACs within a cell [43]. Yeast cells maintained at 37 °C were passaged in the presence of DMSO (control) or an increasing dose of SAHA for 2 days and subsequently imaged to qualitatively assess any alteration in yeast morphology (Fig. 1B). Interestingly, inhibiting all classic HDAC activity with SAHA disrupted normal yeast growth at 37 °C and resulted in improper filamentation in a dose-dependent manner (Fig. 1C). To determine if this phenotype could be attributed to a narrower subset of HDACs in the cell, we assessed the effects of class-specific HDAC inhibitors. The Class I-specific inhibitors Entinostat and Mocetinostat had a similar effect to SAHA at comparable doses, while Class II inhibitors failed to elicit any effect on cell morphology at 37 °C (Fig. 1D, 1E). The effect of Entinostat was not dependent on the presence of N-Acetyl glucosamine (GlcNAc) in the media (Fig. 1F), which has been shown to stimulate filamentation in *Histoplasma* [13]. Entinostat also hastened the onset of filamentation in cells transitioned to growth at RT (Fig. 1G) in a similar manner to what has been observed with cAMP-driven filamentation [19].

Because temperature-mediated morphogenesis is tightly conserved across *Histoplasma* species [18], we queried the effect of Entinostat in strains that are distinct from our North American laboratory strain, G217B. Entinostat disrupted yeast morphology and induced filamentation at 37 °C in *Histoplasma* strains isolated from Central America and Africa (Fig. 1H). This shared effect suggests a conserved requirement for Class I HDAC activity in temperature-mediated morphogenesis of *Histoplasma* at 37 °C.

### Chemical inhibition of Class I Histone Deacetylases disrupts temperature-dependent transcription in *Histoplasma*

Prior transcriptomics studies have identified and characterized unique gene expression signatures that underlie each morphological state in *Histoplasma* [11–13, 44]. Given the morphology phenotypes we observed following Class I HDAC inhibition, we next determined if inhibition of Class I HDACs disrupted the normal gene expression program at 37 °C. We used transcriptional profiling to assess the gene expression signatures of cells undergoing a chemical (HDACi) or physiological (ambient temperature) cue driving filamentation. We grew cultures for 1-2 days at 37 °C with either 10 µM or 50 µM of the Class I HDACi Entinostat (Ent). In parallel, we passaged yeast cells with vehicle (DMSO) at 37 °C and room temperature (RT) to serve as comparison points for both morphological states (yeast and early hyphae, respectively) as shown in Fig. 2A. In contrast to untreated cells that grew primarily as chains of elongated cells shortly after the shift to RT, cells treated with 50 µM of Entinostat formed filaments robustly at RT and grew as a mix of hyphae and elongated cells by day 2 (Fig. 2A). We performed RNA-seq and subjected our data to Principal Component Analysis (PCA) to generate a comprehensive view of transcript abundance variance across these conditions (Fig. 2B). Overall, Entinostat-treated cells appeared to be transcriptionally distinct from both untreated yeast and early hyphae, with a more pronounced difference for the 50 µM treatment (Fig. 2B). The second principal component highlighted a shared transcriptional response in early RT-shifted and Entinostat-treated cells that may represent a common set of gene expression changes associated with the onset of filamentation (Fig. 2B).

We performed differential gene expression analysis of control yeast cells at 37 °C vs cells undergoing both filamentation cues to identify the pathways and genes driving the global signatures observed by PCA. While slight temporal and dose-dependent effects across conditions were observed, the expression profiles were highly correlated between the two doses on the same day and between the two days for the same Entinostat treatment, with a slightly stronger effect being observed at day 2 (Fig. S2A, S2B, and S2C). By day 2, 4241 transcripts were differentially expressed (> 2-fold change, 5% FDR) across the three conditions: 2421 following RT shift, 1243 with 10 µM of Entinostat, and 2087 with 50 µM Entinostat (Fig. 2C).

We used UpSet plots [45] to compare differentially expressed gene (DEG) sets defined across the conditions we tested. Unsurprisingly, a vast majority of the 10 µM Entinostat signature was shared with the effect observed at a higher dose (Fig. 2D and 2E). While more than half of the genes repressed by Entinostat were also repressed during early RT shift (Fig. 2D), the number of shared upregulated genes across these two sets was noticeably smaller (Fig. 2E). Conventionally, Class I HDACs are thought to repress transcription [46], so we were interested in identifying the genes encoding transcripts that showed an inappropriate increase in abundance in 50 µM Entinostat as they could be regulatory targets of interest. We expanded our analysis to include DEG sets defined in steady-state hyphae [18] since cells grown in 50 µM of Entinostat morphologically resembled these hyphae more closely than cells undergoing filamentation at RT at these timepoints. These comparisons revealed that a much larger fraction of upregulated genes was shared between Entinostat-treated cells and steady-state hyphae than between RT-shifted cells and steady-state hyphae (Figs. S2D, S2E). A majority of the genes repressed by Entinostat were also repressed under conditions that promote filamentation (Fig. S2F). Furthermore, the correlation between the signatures of Entinostat and stationary hyphae was stronger than that between Entinostat and acutely RT-shifted cells, both globally and with respect to phase-specific genes (Fig. 2F, 2G, and S2D). These comparisons suggest that Entinostat treatment was inducing a program that normally emerges over time as hyphae develop at RT.

The genes whose expression was dysregulated by Class I HDAC inhibition have functions that are critical to the basic biology of *Histoplasma* yeast or hyphae. A majority (22/40 yeast-specific and 103/165 hyphae-specific) of canonically morphology-responsive genes [14] were significantly dysregulated following Class I HDAC inhibition. This set of genes includes secreted effectors required for virulence *in vivo* such as *CBP1*, *YPS3*, and *KNOT1* [5–7, 9–10] (Fig. 2H). These yeast-enriched effectors are strongly repressed in Entinostat-treated cells in a manner comparable to that of cells undergoing shift to RT (Fig. 2H). Additionally, despite Entinostat-treated cells being cultured at 37 °C, we observed induction of the hyphae-associated transcripts *MS8* [47–48] and *MS95/DDR48* [13, 18, 49] analogous to the induction of these genes shortly after the shift to ambient temperature (Fig. 2H). Interestingly, we also observed the improper expression of transcription-factors (TFs) that promote filamentation in *Histoplasma* [13–14] and morphogenesis in related fungi [50–52]. We observed strong induction of *STU1*, *PAC2*, *NOS1*, *ESDC*, and *FBC1* in Entinostat-treated cells that was not observed following early RT-shift (Fig. 2H) but is typically observed at later timepoints in steady-state hyphae [18]. To determine if the ability of Entinostat to trigger the hyphal program was dependent on a subset of regulatory factors, we compared the Entinostat response of WT *Histoplasma* with strains lacking individual TFs. We determined that filamentation induced by Entinostat at 37 °C requires the presence of *FBC1*, a C2H2 family TF required for filamentation at RT, but not *PAC2*, a WOPR family TF dispensable for filamentation at RT [19] or *VEA1*, a Velvet domain TF that reduces the rate of filamentation after shift to RT [53, 107] (Fig. 2I). Altogether, our transcriptional profiling revealed that Class I HDAC inhibition at 37 °C disrupts the expression of key yeast-associated genes while simultaneously promoting the improper expression of hyphae-associated genes, including regulators that promote filamentation. These results suggest a role for Class I HDACs in mediating temperature-dependent gene regulation by maintaining the regulatory networks that promote the expression of yeast-phase genes at 37 °C while simultaneously repressing regulators that promote filamentation.

### The *RPD3* gene is required for yeast-phase growth at 37 °C

Through our phylogenetic analysis shown in Fig. 1A, we identified two Class I HDACs encoded in *Histoplasma* that we herein named *RPD3* and *HOS2*. To determine a genetic basis for the phenotypes observed following Class I HDAC inhibition, we used a CRISPR/Cas9-based system [53] optimized for use in *Histoplasma* to generate deletion mutants for these HDACs in G217B. We transformed wild-type (WT) yeast with plasmids expressing Cas9 alongside a pair of sgRNAs designed to excise a majority of each gene (Fig. 3A). Following transformation, we screened for successful knock-out (KO) mutants via a colony PCR assay (Fig. S3A and S3B) and validated successful KO strains through whole-genome sequencing (Fig. S3C and S3D). The morphology of *rpd3Δ* cells was reminiscent of the effect of Class I HDAC inhibitors in WT cells at 37 °C. At 37 °C, cells lacking *RPD3* formed filaments (Fig. 3B) and wrinkled colonies (Fig. S3E), whereas *hos2Δ* strains displayed no observable colony or cell morphology phenotype (Fig. 3B, and S3E). In addition to having aberrant morphology at 37 °C, *rpd3Δ* cells also displayed a slight growth defect (Fig. S3F) and enhanced susceptibility to cell wall stress caused by Congo Red and CFW (Fig. S3G).

We generated a construct expressing *RPD3* under the *ACT1* promoter rather than the *RPD3* promoter, for technical reasons. The construct was transformed in parallel into WT and *rpd3Δ* strains. Re-introduction of *RPD3* fully rescued the ability of *rpd3Δ* strains to grow as yeast at 37 °C (Fig. 3D, 3E). Additionally, overexpression (OE) of *RPD3* under the control of the *ACT1* promoter in the WT background did not appear to have any adverse effect on morphology (Fig. 3D). To assess if complementation of the *rpd3Δ* phenotype by *RPD3* required its HDAC activity, we used protein alignments to map the conserved histidine residues required for catalytic activity [54] (Fig. S4A) in *Histoplasma* Rpd3 and generated a plasmid expressing a catalytically dead (CD) Rpd3 mutant. Transformation of this plasmid into *rpd3Δ* failed to rescue the morphology phenotype (Fig. S4B) suggesting the HDAC activity of Rpd3 is required for normal yeast growth. However, sequence-based methods to validate maintenance of the CD plasmid across multiple transformations indicated that this plasmid was poorly maintained in *rpd3Δ* in a manner that was not observed in WT strains or for the WT *RPD3* plasmid in the *rpd3Δ* strain. Importantly, these findings reveal that the effect of Entinostat is largely attributable to inhibition of Rpd3 function, and indicate that the activity of Rpd3, but not Hos2, is required to maintain the yeast phase at 37 °C.

### *RPD3* is required for the pathogenic yeast-phase transcriptional program at 37 °C

To investigate the effect of *RPD3* and *HOS2* on the transcriptional landscape, we subjected WT and mutant strains to RNA-seq. Cultures of WT, *rpd3Δ* and *hos2Δ* strains were grown at 37 °C and then passaged in parallel at 37 °C and RT for 2 days. In contrast to the elongated cells produced by WT and *hos2Δ* cells 2 days following growth at RT, *rpd3Δ* cells formed filaments robustly and grew uniformly as hyphae (Fig. 4A). We performed RNA-seq on these samples and through PCA analysis found that nearly half of the variance in transcript levels (PC1, 47.49%) across both conditions was driven by Rpd3 (Fig. 4B). Of note, a portion of this component is shared between WT and *hos2Δ* cells grown at RT and *rpd3Δ* grown at 37 °C, which suggests a subset of the physiological RT transcriptome is improperly induced in the *rpd3Δ* strain.

We compared transcript abundance between WT cells at 37 °C and RT as well as WT vs each mutant at each temperature. We defined 5480 DEGs in this dataset through these comparisons (Fig. 4C). At 37 °C we identified 2618 DEGs in *rpd3Δ* vs WT and 745 in *hos2Δ* vs WT, while at RT we defined 3585 DEGs in *rpd3Δ* vs WT and 2239 in *hos2Δ* vs WT. The robust signal at RT for *rpd3Δ* is consistent with the strong filamentation phenotype observed under these conditions (Fig. 4A). Altogether, these observations encompass changes to over half of the annotated transcriptome of *Histoplasma* [18].

To focus on the underlying gene expression changes driving the *rpd3Δ* phenotype, we subjected the DEGs defined in our 37 °C data to comparative analysis. At 37 °C in *rpd3Δ* cells, the abundance of 1610 transcripts increased whereas the abundance of 1008 transcripts decreased. These data are in contrast to *hos2Δ* yeast at 37 °C where 495 transcripts increased in abundance and 250 decreased in abundance, consistent with the lack of phenotypic changes in the *hos2Δ* mutant. We generated UpSet plots to compare the DEG sets identified in the mutants at 37 °C alongside time-matched WT cells following shift to RT. We found that the largest intersecting set of DEGs in both directions was shared between *rpd3Δ* at 37 °C and WT cells undergoing filamentation at RT (Fig. 4D and 4E). This finding corresponded with our global observations via PCA and, to an extent, recapitulated our findings using Entinostat to disrupt Rpd3 function. A vast majority (85.8%) of phase-specific genes was dysregulated in *rpd3Δ*: 144/165 hyphae-specific transcripts accumulate improperly at 37 °C, and 32/40 yeast-specific transcripts were improperly depleted at 37 °C (Fig. 4F). We compared the global signatures of *rpd3Δ* and Entinostat-treated cells at 37 °C and found that they were strongly correlated (r= 0.76) (Fig. 4F). A majority of differentially expressed genes were also shared amongst the two conditions (Fig. 4H and 4I). We compared the global profiles of our HDAC mutants to steady-state hyphae and found that *rpd3Δ* but not *hos2Δ* transcriptionally mirrored stationary hyphae and dysregulated phase-specific genes in a manner that corresponded with our observations with Entinostat (Fig. S5A, S5B, and S5C). Furthermore, we also found that more DEGs were shared between *rpd3Δ* and steady-state hyphae than between Entinostat-treated cells and steady-state hyphae (Fig. S5D and S5E). When we further interrogated these gene sets, we similarly discovered strong depletion of key yeast-associated transcripts such as *CBP1*, *YPS3*, *LDF1,* and *KNOT1,* and simultaneous accumulation of hyphae-associated transcripts, including *MS8*, *FBC1, STU1*, *YAP1*, and *ESDC* in *rpd3Δ* cells at 37 °C (Fig. 4G). Of note, this effect appeared to be magnified in *rpd3Δ* grown at RT and in a few instances, for *hos2Δ* at RT (Fig. 4G). Altogether, we uncovered large-scale disruptions to gene expression shared between *rpd3Δ* and Entinostat-treated cells at 37 °C that encompassed much of the canonical temperature-regulated gene network. These findings further support a role for Rpd3 in maintaining the normal regulatory landscape in *Histoplasma* that underlies yeast-phase growth in response to temperature.

### Rpd3 regulates the activity of transcription factors that drive thermal dimorphism

Virtually all yeast transcripts (96%) depend on a network of temperature-responsive transcription factors (Ryp1-4) for their proper expression at 37 °C [15–17]. The Ryp TFs bind multiple sites in the genome and share a number of direct regulatory targets [17]. While a majority of Ryp-mediated gene regulation in yeast appears to be indirect, the set of direct targets shared by Ryp1-3 is enriched for yeast-specific genes [17, 55]. We noticed that a considerable number of Ryp-dependent genes are transcriptionally dysregulated following chemical and genetic disruption of Rpd3. We compared our DEG sets to the shared targets of Ryp1-3 previously determined by chromatin immunoprecipitation (ChIP) [17] and determined that this core set of yeast-specific genes was significantly enriched for genes that were dysregulated following a high dose of Entinostat or KO of *RPD3* (Fig. 5A). 24.2% of shared Ryp1-3 targets were dysregulated in 50 µM of Entinostat (Fisher’s exact test, p = 0.019) while 38.7% were dysregulated in *rpd3Δ* (Fisher’s exact test, p = 1.66e-06) (Fig. 5A).

To determine if disruption in yeast phase-specific gene expression following KO of Rpd3 could be attributed to a change in Ryp protein levels, we performed Western blotting in WT cells following 2-day exposure to DMSO or 50 µM Entinostat and *rpd3Δ* cells at 37 °C. We detected comparable levels of all three Ryp proteins between the two *RPD3* perturbations and their controls with a negligible decrease in Ryp1 and Ryp3 in *rpd3Δ* (Fig. 5B), indicating that changes in the transcriptome were not due to depletion of Ryp transcription factors.

Rpd3 has been found to influence the binding and activity of stress responsive TFs that regulate the response to heat shock in *S. cerevisiae* [27–30]. Because Ryp-dependent genes are dysregulated following Rpd3 disruption, we hypothesized that Rpd3 might affect the ability of the Ryp TFs to associate upstream of their regulatory targets. We performed ChIP-seq to investigate the genome-wide binding patterns of Ryp1-3 in WT cells following 2 days of growth with DMSO or 50 µM Entinostat at 37 °C. Ryp signal in control cells was strong at the *CBP1* locus, recapitulating our previous observations (Fig. 5C) [17]. Strikingly, this signal was predominantly lost in cells treated with Entinostat (Fig. 5C), indicating that Entinostat affected Ryp association with the DNA. We similarly observed a decrease in Ryp ChIP signal upstream of *CBP1* in *rpd3Δ* cells in comparison to WT cells (Fig. 5D). We used the MACS2 peak finding algorithm [89] to identify sites of Ryp enrichment in promoters across the genome. Although the Ryp2 peak signal was muted, making the Ryp2 analysis difficult, we identified high-quality peaks for Ryp1 and Ryp3 in both control and treatment conditions. We found that the vast majority of Ryp1 and Ryp3 binding events were lost in cells grown with Entinostat (Fig. 5E). We identified 102 Ryp1 and 236 Ryp3 binding events in control cells, whereas only 7 Ryp1 and 46 Ryp3 peaks were identified in Entinostat-treated cells (Fig. 5E). Of these peaks, only 2 Ryp1 peaks and 20 Ryp3 peaks were shared between the two conditions (Fig. 5E). These findings suggest class I HDAC function is required to maintain the normal genome-wide binding patterns of the Ryp TFs that takes place at 37 °C.

We then assessed differential Ryp binding by quantifying and normalizing ChIP signal in MACS2-defined [56] Ryp binding sites and generated contrasts between control and Entinostat-treated cells. We observed a comprehensive decrease in Ryp ChIP signal in peaks upstream of known and putative effector genes [10] in cells undergoing treatment with Entinostat (Fig. 5F). We identified Ryp peaks in *rpd3Δ* strains using MACS2 and similarly found that a majority of Ryp1-3 binding events were lost in *rpd3Δ.* In total, 617/775 of the Ryp1 peaks, 266/406 Ryp2 peaks, and 429/658 of the Ryp3 peaks identified in WT cells were lost in *rpd3Δ* (Fig. 5G, 5H, and 5I). Notably, we also identified a substantial number of Ryp1-3 targets that were unique to *rpd3Δ*. We identified 244 Ryp1, 216 Ryp2, and 243 Ryp3 bound genes in *rpd3Δ* strains that were not present in WT cells (Fig. 5G, 5H, and 5I). This suggested that loss of *RPD3* may drive changes to chromatin structure that result in significant changes to association sites of the Ryp proteins across the genome. We followed our peak-level analysis with quantification of differential ChIP signal in WT-defined peak loci in *rpd3Δ* and similarly observed a loss of Ryp signal in the promoters of putative effector genes that are typically enriched in yeast phase (Fig. 5J). Combined, these findings point to a role for Rpd3 activity in maintaining the normal regulatory landscape that facilitates Ryp binding and activity at 37 °C.

### Rpd3 disruption alters histone acetylation patterns at the loci of known and putative regulators of filamentation

Rpd3 regulates responses to thermal and osmotic stress in *S. cerevisiae* by targeting histones for deacetylation at the loci of stress-responsive genes [28–29]. This hypoacetylation subsequently decreases genome accessibility to restrict the binding of stress-responsive TFs and blocks the recruitment of transcriptional machinery [28–29]. At the time of our study, no experimental datasets had defined or characterized histone acetylation patterns in *Histoplasma,* so we based our selection of Rpd3 substrates on findings in other fungi. In *S. cerevisiae,* Rpd3 is reported to regulate transcription through the removal of acetyl groups deposited on N-terminal lysine residues of histones H3 and H4 [46, 54]. We used ChIP-seq to globally query the dynamics of histone H3 acetylation following chemical or genetic inhibition of Rpd3 at 37 °C. We performed pull downs on acetylated H3K9 (H3K9Ac) and H3K14 (H3K14Ac), two marks that label transcriptionally active loci [46] targeted by Rpd3 for deacetylation and subsequent transcriptional silencing [57]. We hypothesized the loci surrounding genes normally deacetylated (and thereby transcriptionally repressed) by Rpd3 at 37 °C may display signs of hyperacetylation when Rpd3 function is compromised. We generated coverage tracks of normalized histone acetyl-ChIP signal and found that loci encoding TFs with putative roles in filamentation are hyperacetylated following disruption of Rpd3. Notably, these TFs were also found to be transcriptionally induced following either chemical or genetic perturbation of Rpd3 (Fig. S6A). One of the transcripts most strongly upregulated following either Rpd3 disruption or in steady-state hyphae is a homolog of *esdC* in *Aspergillus spp.*, a putative regulator of sexual development [52]. We observed increased H3K9Ac and H3K14Ac signal spanning the *ESDC* promoter and gene body in Entinostat-treated and *rpd3Δ* cells (Fig. 6A) that was not present in control cells. A homolog of *YAP1*, a bZIP TF that regulates environmental stress responses in *S. cerevisiae* [58], displayed similar hyperacetylation of H3K9 and H3K14 across its promoter and gene body regions (Fig. 6B) and underwent transcriptional induction following either perturbation of Rpd3 (Fig. S6A). A similar result was observed surrounding the start site of *I7I48_01662*, a hybrid homeobox/C2H2 zinc-finger TF identified as being strongly induced in conditions that drive filamentation [19] (Fig. 6C).

We also observed changes in acetylation patterns that correspond with two alternative transcript start sites (TSS) in the 5’ region of *WET1*, a TF whose regulation is critical to *Histoplasma* morphology [18]. These alternate TSSs yield *WET1* transcripts with two distinct leader sequences- a longer leader in yeast that is poorly translated, and a shorter leader in hyphae that is robustly translated [18]. We observed strong H3K9Ac acetylation peaks surrounding the yeast-specific TSS of *WET1* in control cells that was lost in Entinostat-treated and *rpd3Δ* cells (Fig. 6D). Simultaneously, we also observed an increase in H3K9 acetylation surrounding the downstream hyphae-specific TSS and spanning the hyphal isoform of the *WET1* transcript (Fig. 6D). Of note, we also observed instances of Rpd3-independent H3K9 hypoacetylation at the loci of yeast-specific genes (Fig. S6B and S6C), suggesting that there may be additional HDAC/HAT complexes that regulate chromatin state to silence yeast-specific genes following the onset of filamentation.

We quantified differential H3K9Ac and H4K13Ac ChIP signal on ATG-defined promoter regions across the *Histoplasma* genome to globally assess changes to histone acetylation. Overall, only a small subset of promoters displayed signs of differential H3K9Ac and H3K14Ac signal following Entinostat treatment and *RPD3* disruption. Similar pulldowns and promoter analysis of acetylated H4K16, a modification that is not considered a primary regulatory target of Rpd3 [60], failed to identify instances of differential promoter acetylation across the entire genome (Table S12). We indexed our differential promoter signal generated against H3K9Ac and H3K14Ac on putative TF-encoding genes across the *Histoplasma* genome (Fig. 6E). Clustering this set of data highlighted a subset of TFs with hyperacetylated promoters following Entinostat treatment or KO of *RPD3* (Fig. 6E). This quantification corroborated our manual observations of ChIP-seq coverage in the promoters of *ESDC*, *WET1*, and *YAP1* (Fig. 6F). We additionally observed increased acetylation signal in the promoters of the known and putative pro-filamentation TFs such as *STU1*, *VELC*, *NOS1* and *WET1* (Fig. 6F). Notably, these TFs were also among those we identified for being transcriptionally induced following chemical or genetic targeting of Rpd3 (Fig. S6A). These findings suggest Rpd3 may function to drive deacetylation at the loci and cis-regulatory regions of pro-filamentation TFs to repress their inappropriate transcription in yeast cells at 37 °C.

### Rpd3 is required for virulence in a macrophage model of infection

As Rpd3 regulates a majority of defined temperature-specific genes (Fig. 4F), we hypothesized that its regulon would encompass virulence genes coupled to yeast-phase growth. We indexed our RNA-seq data following chemical and genetic ablation of Class I HDACs on a comprehensive set of putative effectors across the G217B genome previously defined based on a combination of high throughput experiments and sequence features [10]. We found that most of the putative effectors encoded in G217B are repressed at 37 °C when Rpd3 function is chemically or genetically compromised (Fig. 7A). Similarly, these transcripts show decreased abundance in control cells undergoing filamentation at RT (Fig. 7A).

**Figure 7.**
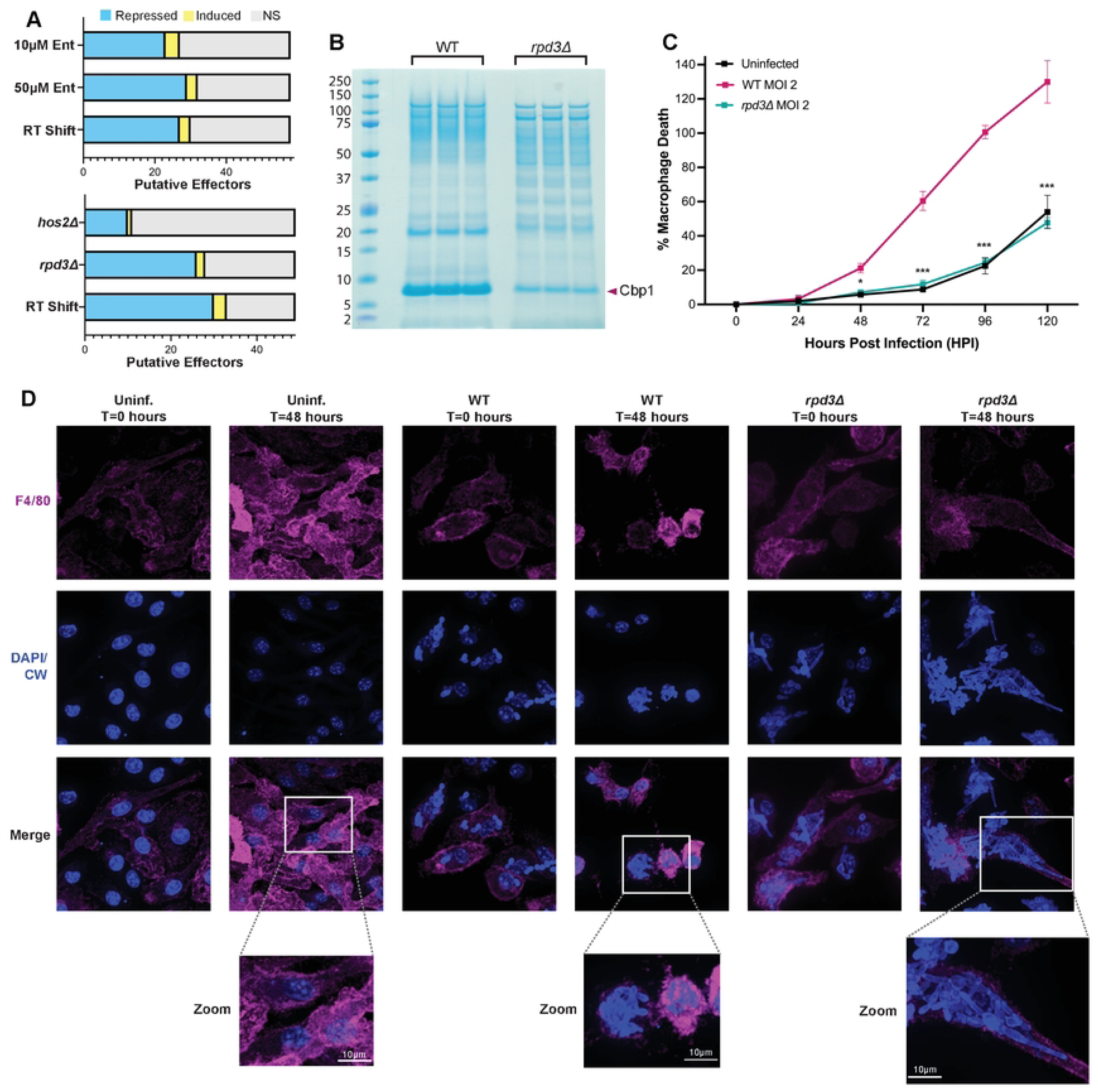
*rpd3Δ* lack the expression of key virulence genes and fail to lyse macrophages. A. Stacked boxplots listing the number of putative virulence effectors encoded in G217B (defined by Rodriguez *et al.* 2025 [18]) that are transcriptionally dysregulated following early RT shift and chemical (10 µM and 50 µM Entinostat) or genetic (*rpd3Δ* and *hos2Δ*) disruption of Class I HDACs. These plots represent a subset of the differential genes identified in Figs. 2C and 4C, respectively. B. SDS-PAGE gel of culture supernatants from WT and *rpd3Δ* strains grown to saturation at 37 °C stained with Instant Blue. The arrowhead on the right indicates Cbp1 signal and protein standard molecular weights (in kDa) are indicated on the left. C. Lactate dehydrogenase (LDH) release assay used to quantify the lysis of bone barrow-derived macrophages (BMDMs) (as % Macrophage Death) over the time-course of infection. BMDMs were mock-infected (uninfected) or infected with G217B WT and *rpd3Δ* at an MOI of 2 and assayed for LDH release every 24 hours for 5 days. Asterisks indicate statistical significance from t-tests comparing WT and *rpd3Δ-*infected BMDMs at each timepoint: * = p < 0.5, *** = p < 0.001 D. Confocal fluorescence microscopy images of mock-infected (uninf.) or G217B WT and *rpd3Δ-*infected BMDMs at an MOI of 2 captured at day 0 (T=0) and day 2 (T=48 hours) post-infection. Samples were fixed at each timepoint and incubated with a fluorophore-conjugated F4/80 antibody before being stained with Calcofluor White and DAPI for imaging.

One of the Rpd3-dependent effectors we identified is *CBP1*, a secreted virulence factor required for macrophage lysis *in vitro* and pathogenesis in a mouse model of infection [5–9]. *CBP1* is one of the most abundant and efficiently translated transcripts produced by *Histoplasma* yeasts [18], so we speculated that the decrease in Cbp1 transcript abundance observed when Rpd3 is compromised would be sufficient to result in a decrease to Cbp1 protein levels. We used an established SDS-PAGE assay on concentrated culture supernatants [9–10] harvested from WT and *rpd3Δ* cultures grown to saturation at 37 °C to qualitatively assess Cbp1 protein levels. We observed a marked decrease in the abundance of Cbp1 in *rpd3Δ* strains in addition to other apparent alternations in the profile of secreted proteins (Fig. 7B). We previously found that the extent of macrophage lysis following infection with *Histoplasma* correlates with the abundance of Cbp1 [8]. To determine if the repression of virulence genes, including Cbp1, in *rpd3Δ* affects *Histoplasma* virulence in a macrophage model of infection, we infected murine bone-marrow derived macrophages (BMDMs) with WT or *rpd3Δ Histoplasma* and measured macrophage lysis as previously described [7, 99]. We found that the *rpd3Δ* mutant is completely unable to lyse BMDMs and that *rpd3Δ*-infected BMDMs are virtually indistinguishable from uninfected BMDMs throughout the time-course of infection (Fig. 7C). This was in stark contrast to WT-infected BMDMs that were entirely lysed by the end of the infection (Fig. 7C).

*Histoplasma* yeasts attain a high intracellular fungal burden prior to the onset of macrophage lysis [61]. We wanted to verify that the striking lysis defect in *rpd3Δ* was not confounded by an intracellular growth defect in BMDMs as the *rpd3Δ* mutant displayed a slight *in vitro* growth defect (Fig. S2F). Due to its altered morphology, we faced technical challenges with plating-based methods of assessing intracellular growth of *rpd3Δ* so, as an alternative, we subjected BMDMs infected with WT and *rpd3Δ* to fluorescence imaging to qualitatively assess intracellular growth over the time-course of infection. Remarkably, we observed that *rpd3Δ* cells replicated within macrophages and reached a high fungal burden without lysing BMDMs (Fig. 7D). Interestingly, this recapitulates the phenotypes of strains lacking a functional copy of *CBP1*, as *cbp1Δ* yeast are capable of replicating within macrophages but fail to induce host cell lysis [5–9]. In addition, we found that the growth of hyphae that occurs during infections with the *rpd3Δ* strain is not sufficient to mechanically breach the macrophage plasma membrane (Fig. 7D) in contrast to what has been observed in other intracellular fungi that utilize filamentation and polarized growth to induce host cell lysis [62]. These observations indicate that Rpd3 is required for virulence in macrophage infections through its role in promoting the expression of key yeast-associated virulence genes at 37 °C, including Cbp1.

## Discussion

Microbial pathogens utilize complex molecular machinery to sense and adapt to constantly changing environmental conditions. *Histoplasma spp.* and other thermally dimorphic fungi are adept at sensing temperature to trigger gene regulatory responses that enable their survival and persistence across two vastly different niches—the environment and mammalian hosts. When recapitulating the transition in vitro, depending on culture conditions and the direction of the morphologic switch, it occurs over a few days to a few weeks [13–14, 20]. The highly dynamic and prolonged nature of the transition suggests a multifaceted regulatory system that allows *Histoplasma* to distinguish between transient and sustained change in temperature. In this study, we investigated how modifications to chromatin state, a previously unexplored regulatory mechanism in *Histoplasma*, contributes to regulating the cell-fate decisions that maintain the yeast state in response to host temperature. We identified the class I HDAC Rpd3 as required for yeast-phase growth and virulence factor expression under host-like conditions. When Rpd3 function was compromised, either chemically or genetically, the cells could no longer maintain the appropriate morphologic and molecular response to temperature, and instead defaulted to a hyphal growth program even at 37 °C. While we found that the remaining class I HDAC in *Histoplasma,* Hos2, does not play a critical role in regulating *Histoplasma* development in response to temperature, it plays significant roles in morphogenesis, virulence, and gene expression in entomopathogenic and phytopathogenic fungi, including the thermally dimorphic plant pathogen *U. maydis* [35–36]. Despite there being functional divergence between Hos2 and Rpd3 homologs across fungi, the combined contributions of class I HDACs in virulence and morphotype switching remain highly conserved across different evolutionary clades within the fungal kingdom [39, 42], highlighting the importance of chromatin remodeling in key aspects of fungal development.

With respect to morphology and gene expression, short-term disruption to Rpd3 function with Entinostat produces cells that more closely resemble stationary hyphae (normally produced after long-term transitions to RT) than cells that result from short-term transition of yeast cells to RT (Fig. 2A, 2G, 4F, and S2D). The short-term timepoint likely represents a transient snapshot within the transition to mature hyphae. More than 80% of the differential signal in early RT-shifted cells was made up of yeast-associated transcripts that were depleted after shift to RT, presumably to shut off the yeast program prior to the onset of filamentation. The much smaller subset of hyphal transcripts that were being induced did not include key regulators that are sufficient to drive filamentation like *FBC1, PAC2*, and *WET1,* suggesting that the hyphal network has yet to be fully activated at this early timepoint. These findings are in contrast to Entinostat treatment, where close to 60% of the differential signal is composed of transcripts that improperly accumulate at 37 °C—a large fraction of which is typically associated with stationary hyphae (Fig. S2E), including the TFs listed above.

As HDAC activity is normally associated with transcriptional repression, a simple model would predict that disrupting HDAC function triggers gene induction. In our RNA-seq data, we found that that this was not entirely the case since 35-40% of the Entinostat and *rpd3Δ*-induced signatures were composed of transcripts that were decreasing in abundance. This finding isn’t entirely surprising, however, as Rpd3 has been shown to both positively and negatively regulate transcription through influencing the activity of TFs and signaling molecules that both activate and repress the expression of their regulatory targets [27–28, 63]. While the set of transcripts that are depleted following Rpd3 disruption is not fully reflective of the entire repertoire of yeast-specific genes that are normally repressed early following the shift to RT, this set of genes is enriched for direct Ryp1-3 targets previously identified through ChIP studies (Fig. 8) [17]. *S. cerevisiae* Rpd3 is recruited to heat shock genes in response to thermal stress, where its HDAC activity promotes the binding and activity of the Msn2/4 TFs [27–26]. In *C. albicans*, Rpd3 and its paralog Rpd31 were also found to regulate TF activity of the *RYP1* homolog *WOR1* to influence morphogenesis in response to host-associated cues [31]. The enrichment of direct Ryp1-3 targets in the set of Rpd3-dependent genes implies that Rpd3 may similarly influence genomic architecture to direct TF binding in *Histoplasma.* Through our ChIP studies, we found that Ryp activity in the promoters of yeast-associated virulence genes like Cbp1 is dependent on Rpd3 activity. Overall, upwards of 90% of Ryp-binding events across the genome are lost with Entinostat, and 65-80% of Ryp-binding events are lost in *rpd3Δ* (Fig. 5G-I). Thus, Rpd3 in *Histoplasma* may maintain a genomic landscape that facilitates the recruitment of transcriptional machinery and promotes the activity and complex formation of temperature-responsive TFs like Ryp1-3 (Fig. 8).

**Figure 8.**
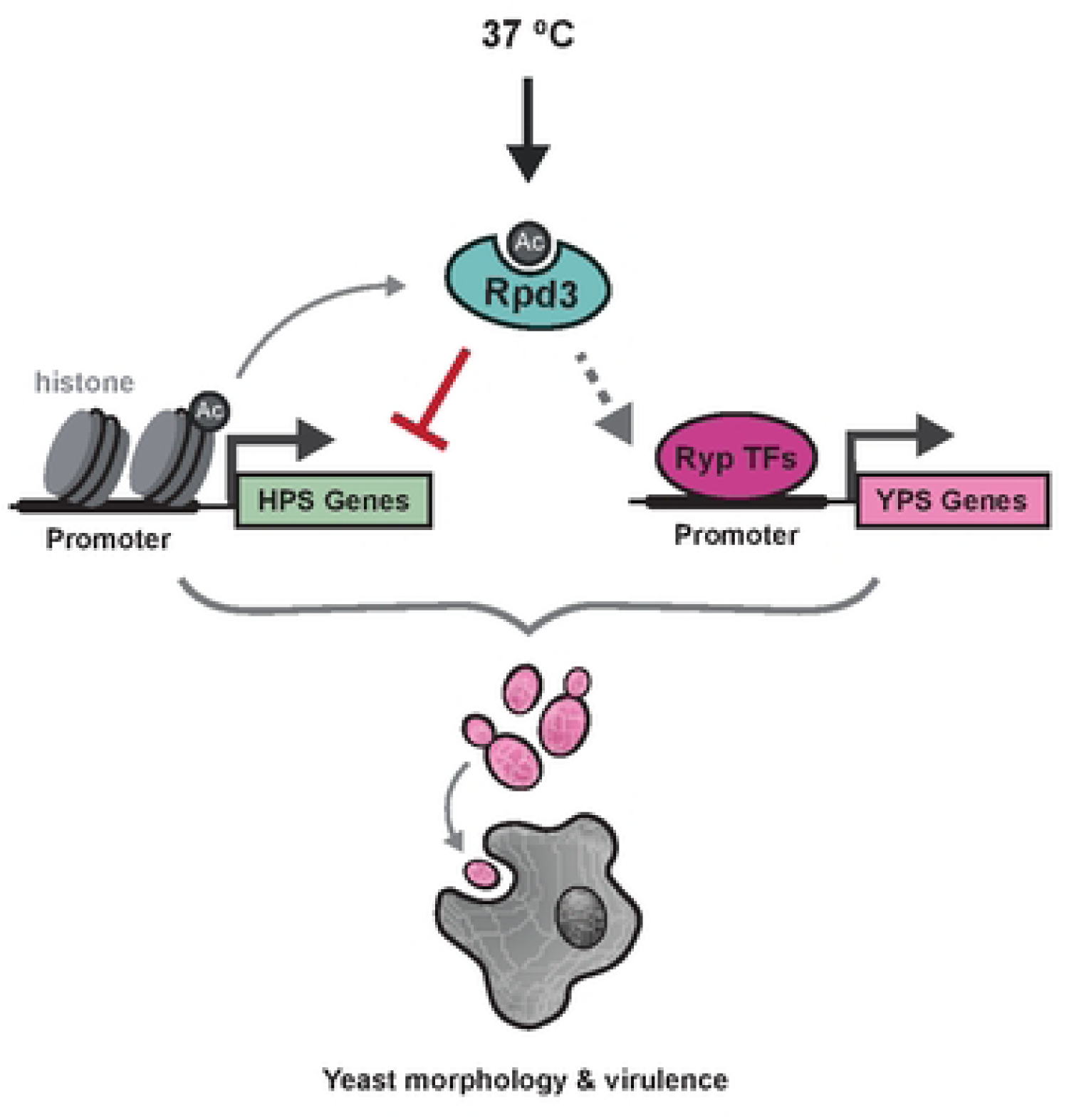
Rpd3 integrates with transcription factor networks to promote *Histoplasma* yeast-phase growth under host conditions. Proposed model for Rpd3 function under host conditions. Rpd3 is required for the repression of hyphal-phase specific (HPS) genes and the expression of yeast-phase specific (YPS) genes at 37 °C. Our data suggest a model where Rpd3 represses HPS gene expression through the removal of acetyl (Ac) groups on histone H3 (histone), most notably in the promoters of HPS transcription factors (TFs) that may antagonize the Ryp network. We hypothesize the genomic landscape established by Rpd3 is indirectly (dashed arrows) required to promote activation of YPS genes through promoting the DNA-binding activity of the Ryp TFs (Ryp1-3). Thus Rpd3 is required for *Histoplasma* yeast-phase morphology and virulence gene expression to promote macrophage lysis at 37 °C.

Another method by which chromatin remodeling factors influence gene activation is through modulating the accessibility of alternative TSSs that fall upstream of an open reading frame (ORF). Prior studies in our lab found that a subset of phase-specific transcripts display alternative TSSs in each phase, thereby creating differential 5’ untranslated regions (UTRs) or leader sequences [18]. In *S. cerevisiae*, Rpd3 regulates the accessibility of different 5’ UTR isoforms upstream of meiotic genes to repress their translation during mitotic growth [64]. We observed Rpd3-dependent changes to acetylation patterns in the 5’ UTRs of *WET1* (Fig. 6D) and MS95 (Fig. S7B) associated with two transcript isoforms that are differentially translated and expressed in yeast and hyphae [18]. The acetylation signal in cells undergoing filamentation following Rpd3 disruption corresponds with a shorter leader sequence that is normally enriched in hyphae and confers robust translation. Rpd3 disruption also results in the loss of hyperacetylation surrounding the yeast-associated TSS sites associated with these genes at 37 °C. We observed additional instances of Rpd3-dependent hyperacetylation at 37 °C upstream of yeast-specific genes (Fig. S6B and S6C) that suggests Rpd3 influences the activity of other histone modifying enzymes to regulate the overall genomic landscape in yeast cell. These findings present a mechanism by which chromatin remodelers like Rpd3 influence the expression of phase specific genes in *Histoplasma* through remodeling the chromatin landscape surrounding alternative TSS sites in response to temperature.

This study provides the first experimental assessment of genome-wide histone acetylation patterns in *Histoplasma.* We found that histone H3 acetylation in the loci surrounding and spanning open reading frames was highly dynamic in the *Histoplasma* genome. While compensation by other HDACs in the cell may have contributed to the lack of differential histone H4 acetylation on H4K16 (Fig. S7) and H4K5 (data not shown), lysine residues on histone H3 may be preferential substrates for Rpd3 in *Histoplasma*. We determined that some pro-filamentation TFs previously characterized by our lab including *FBC1*, *STU1*, and *WET1* display Rpd3-dependent H3K9/14 hypoacetylation that corresponds with their transcriptional repression at 37 °C. In addition to these TFs, we also observed similar instances of Rpd3-dependent hypoacetylation surrounding genes encoding regulators like *YAP1*, *ESDC*, *I7I48_01662*, *NOS1*, and *WHC1.* These TFs are strong candidates for follow-up studies to identify novel regulators of filamentation as they may represent key players in the regulatory networks that indirectly or directly antagonize the Ryp network to repress yeast growth at RT. We hypothesize Rpd3 recruitment to these loci occurs in a temperature-dependent manner to restrict their expression at 37 °C and promote their activation at RT. Rpd3 functions within multiprotein HDAC complexes that include subunits whose interactions with TFs can facilitate the temporal and site-specific localization of Rpd3 across the genome [31].

Prior to its discovery as a potent pan-HDACi, a naturally derived compound, trichostatin A (TSA) isolated from the bacteria *Streptomyces platensis*, was identified for its fungicidal activity against *Aspergillus* and *Tricophyton* [68–69]. While only moderately effective when used on their own, HDAC inhibitors have robust fungicidal activity against a broad panel of human fungal pathogens when used in conjunction with azole and echinocandin antifungals [71–72]. Antifungal resistance is a pressing clinical concern, so these findings point to promising potential for HDAC-targeting combinatorial therapies against fungal infections, especially with the development of fungal-specific pan-and class II-specific HDAC inhibitors [73–74]. The macrophage lysis defect of the *rpd3Δ* strain (Fig. 7C) suggests that targeting Rpd3 in *Histoplasma* may have therapeutic potential. While there has yet to be success in developing a fungal-specific class I HDAC inhibitor, Rpd3 homologs in filamentous fungi harbor a unique extension of their C-terminal regions that is demonstrated to be critical for nuclear localization and enzymatic activity in *Aspergillus* [75] and *Magnaporthe* [38]. This unique feature with functional significance may present as a promising target for the development of novel antifungals with specificity to Rpd3 in filamentous fungi. WT *Histoplasma* grow exclusively as yeast within macrophages, where they trigger host-cell lysis, but the *rpd3Δ* mutant strain exhibited altered intracellular morphologies including filamentous projections. Some intracellular fungal pathogens use filamentation to escape from the restrictive environment within phagocytic immune cells. While filamentation plays key roles in *C. albicans* virulence and facilitates intracellular escape, it is not required for macrophage lysis, which also requires the production of a pore-forming toxin and the induction of pyroptosis through caspase-1 [78–79]. In *Histoplasma,* macrophage lysis relies on secreted yeast-specific effectors like Cbp1 that access the host cytosol and activate the integrated stress response to trigger macrophage cell death [7–9]. Cbp1-triggered host apoptosis ultimately facilitates pathogen escape and the progression of infection [7–8]. In contrast to other intracellular fungi, the mechanical stress induced by intracellular growth of *Histoplasma* yeasts (such as the *cbp1Δ* mutant) or filaments (as we have observed in this study with *rpd3Δ*), is not sufficient to induce macrophage lysis. Our transcriptional data suggest that the lysis defect is a direct consequence of disrupting the expression of effectors like *CBP1*, *YPS3*, and *LDF1*, explaining why *RPD3* is critical for the *Histoplasma* yeast lifestyle.

HDACs are reported to be involved in spore development in related fungi, including *M. oryzae* and the thermal dimorph *U. maydis* [35, 38]. Despite numerous attempts, the *rpd3Δ* mutant did not grow under different sporulation conditions and thus did not generate conidia. This phenotype was unexpected, as a number of conidia-specific genes, including the most highly enriched conidial transcript in *Histoplasma,* GAD1 [12], were strongly upregulated following Rpd3 disruption (Fig. 2H and 4G). We also observed inappropriate H3K9/14 hyperacetylation at the *GAD1* locus and promoter at 37 °C following Rpd3 disruption (Fig. S7C) indicating that Rpd3 may normally restrict accessibility and transcription of this gene under conditions that do not promote conidiation. The hypersensitivity to cell-wall stress displayed by *rpd3Δ* (Fig. S3G) may contribute to its poor viability on sporulation media, so further work to optimize the growth conditions of *rpd3Δ* at RT and under sporulation conditions will be crucial to fully characterize any contributions Rpd3 may play in *Histoplasma* development at ambient temperature.

Epigenetic mechanisms to direct cell fate have long been appreciated in metazoans, however their contributions to the regulatory landscape are less appreciated within the fungal kingdom. The results outlined in this study highlight the functional significance of a histone modifying enzyme in the biology of a human fungal pathogen. Our findings also contribute to a more comprehensive understanding of the *Histoplasma* factors involved in the complex regulatory networks that direct the transcriptional response to temperature (Fig. 8). In addition to Rpd3, our data suggest that other histone-modifying enzymes may contribute to regulating temperature-dependent gene expression. Future studies to assess the contributions of HATs that may functionally oppose HDACs like Rpd3, in addition to histone methyltransferases and demethylases that can influence HDAC/HAT recruitment [42], will be crucial to determine the full repertoire of histone-modifying enzymes that contribute to regulating thermal dimorphism in this ubiquitous pathogen of humans.

## Acknowledgements

We thank members of the Sil and Noble labs for comments on the manuscript. We acknowledge the UCSF PCAT for use of equipment, and the UCSF CAT and CZI Biohub San Francisco for sequencing resources.

## Supplemental Information

**Table S1.**
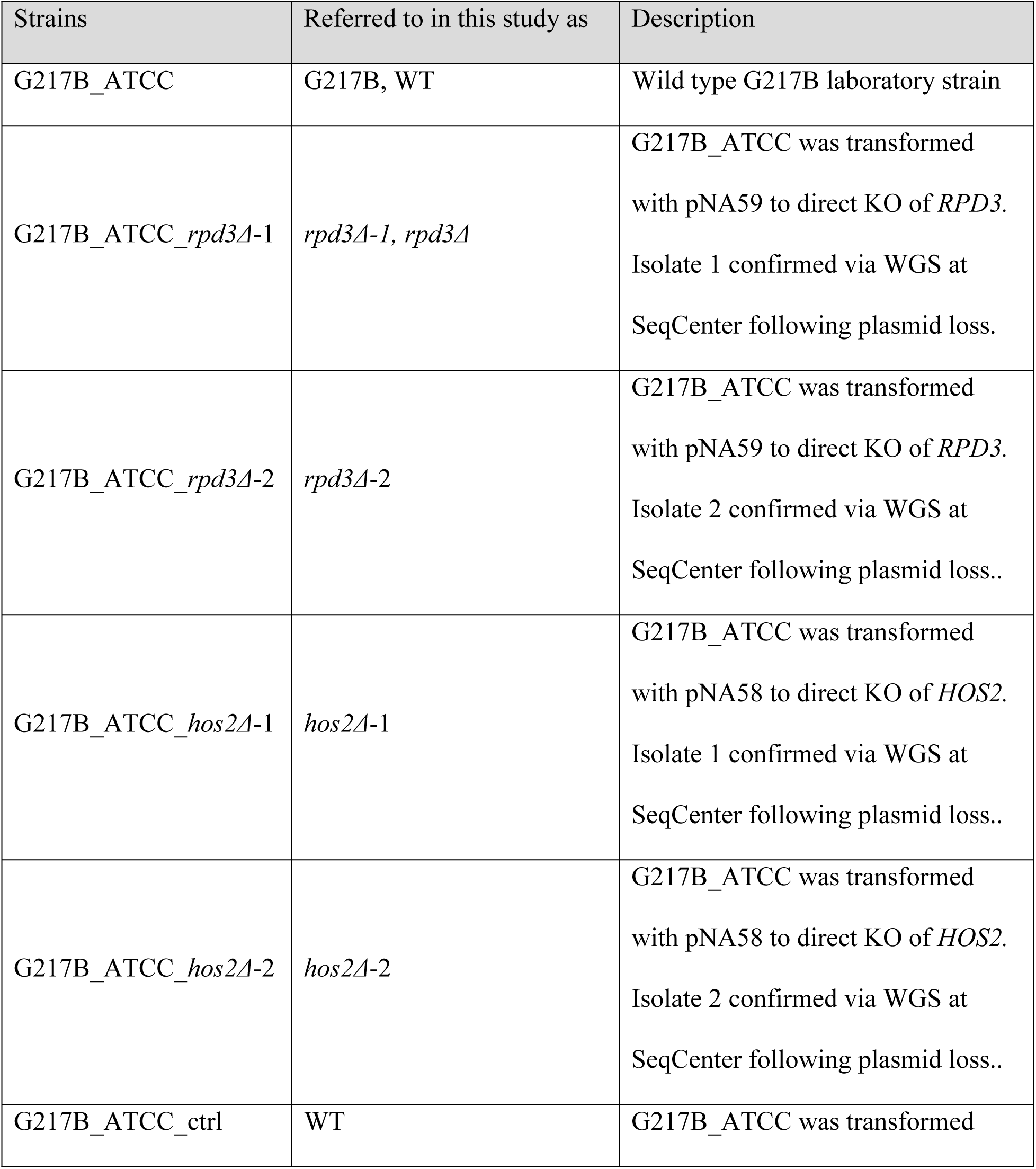
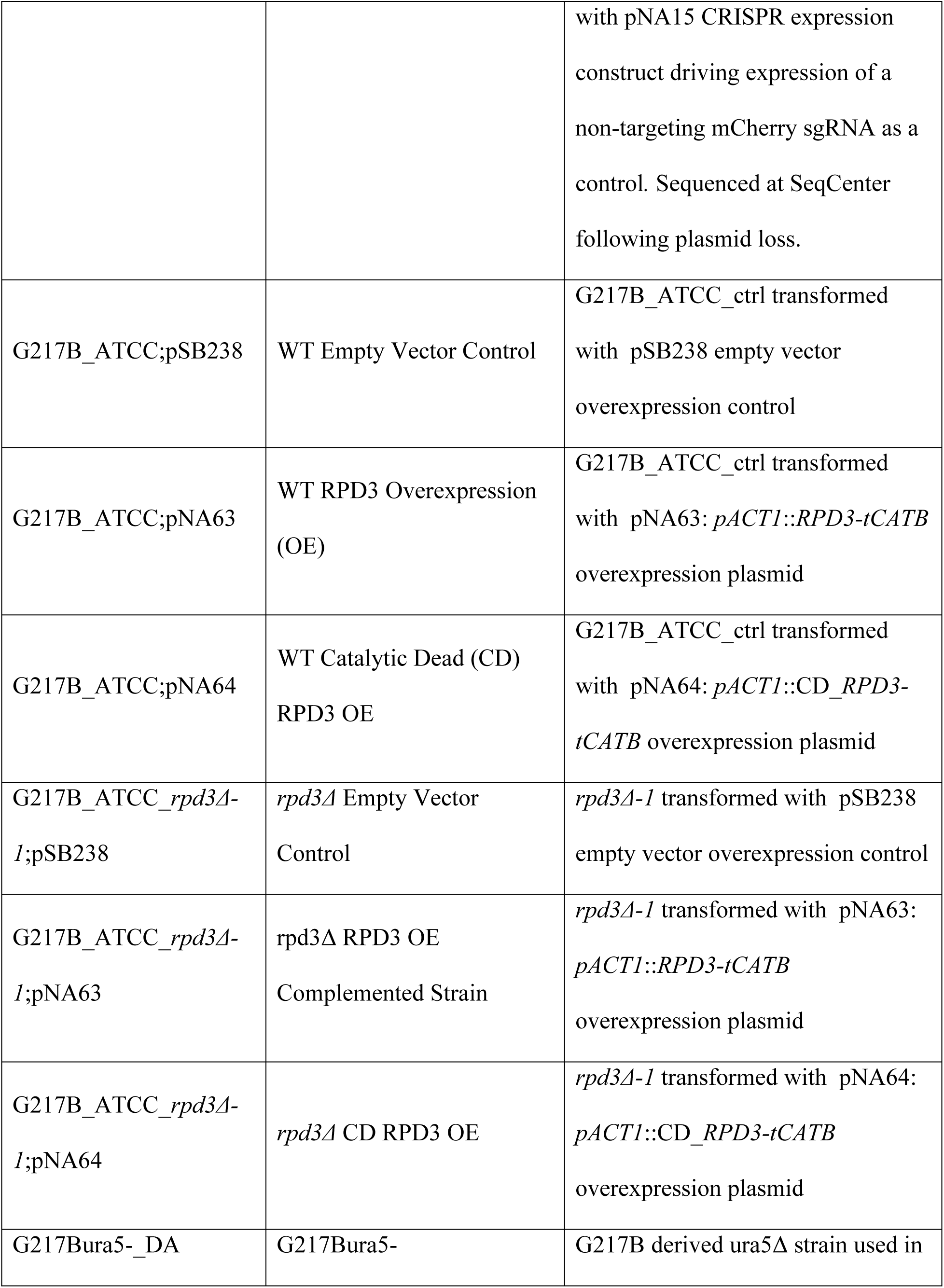
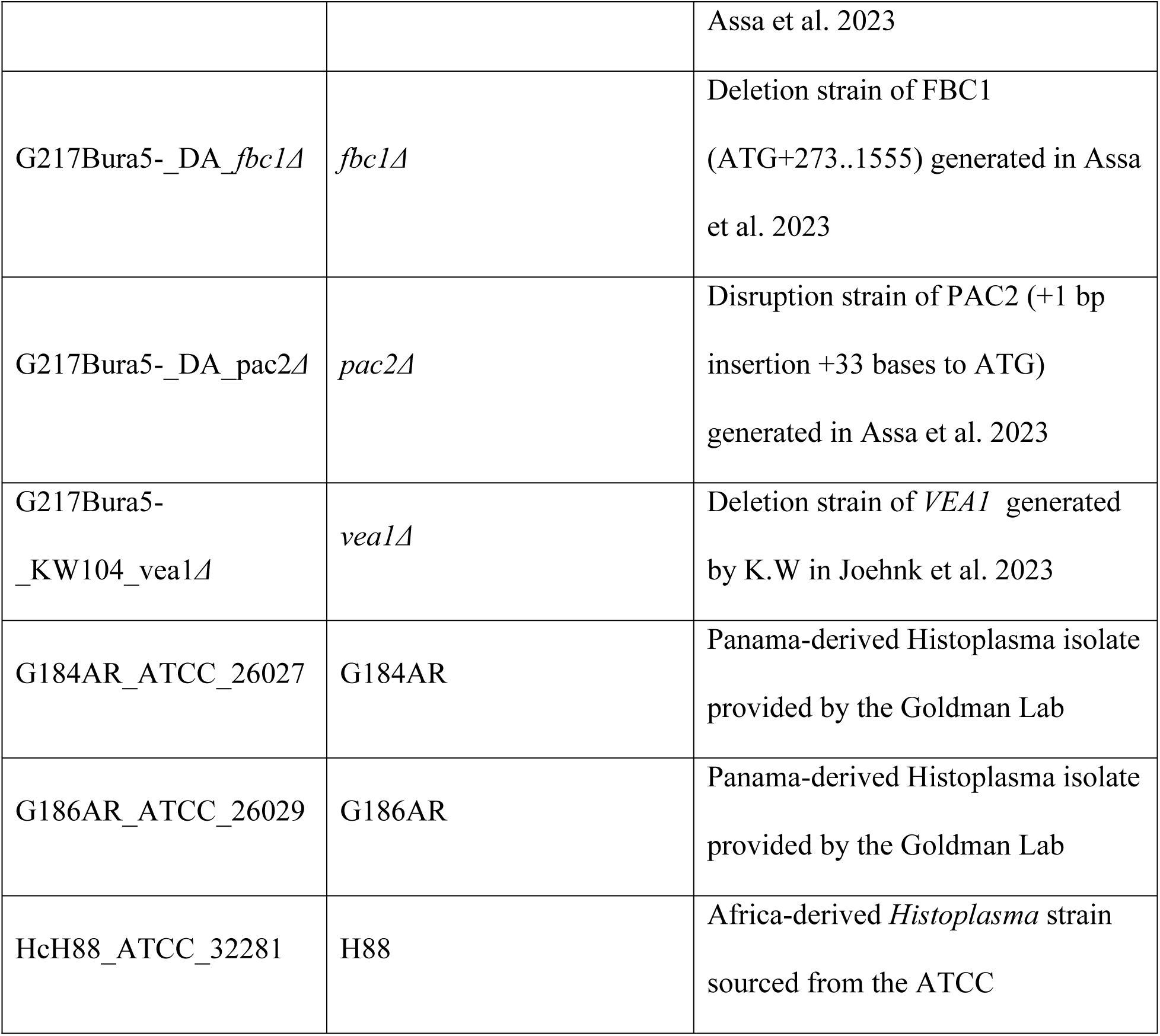
Strains used in this study.

**Table S2.** Plasmids used in this study.

| Plasmid | Description |
| --- | --- |
| pNA15 | *hph* marked mCherry sgRNA-expressing Cas9 plasmid to be used as control plasmid for generating mutant strains with CRISPR/Cas9. |
| pNA16 | pDONR entry clone containing *pGAPDH*-*SrfI*-*ttrpC* used for cloning sgRNA cassettes into via Gibson Assembly following inearization at the *SrfI* site. |
| pNA55 | pDONR entry clone containing *pGAPDH*-gNA14,gNA15-*ttrpC*. gNA14-15 |
|  | represent gBlock sgRNA cassettes with protospacer sequences targeting the 5' and 3' of *HOS2*, respectively. |
| pNA56 | pDONR entry clone containing pGAPDH-gNA16, gNA17-*ttrpC*. gNA16-17 represent gBlock sgRNA cassettes with protospacer sequences targeting the 5' and 3' of *RPD3*, respectively. |
| pNA58 | pNA55 was used for LR with pBJ262 to obtain *hph*-marked dual-sgRNA containing CRISPR/Cas9 expression plasmid for *HOS2* KO. |
| pNA59 | pNA56 was used for LR with pBJ262 to obtain *hph*-marked dual-sgRNA containing CRISPR/Cas9 expression plasmid for *RPD3* KO. |
| pNA61 | pDI71 template amplified with NA139/140 primers to generate a pDONR backbone with pACT1:-empty-tCATB that was assembled with the full *RPD3* ORF amplified from gDNA using NA141/142 to generate the *RPD3* OE cassette entry clone. |
| pNA62 | pNA61 was used for SDM using NA143/144 primers to introduce H150A and H151A mutations in the catalytic domain of *RPD3*. |
| pNA63 | pNA61 was used for LR reaction with pSB238 in order to introduce Ryp3 OE cassette into *hph* marked expression vector used for transformation. |
| pNA64 | pNA62 was used for LR reaction with pSB238 in order to introduce *RPD3* H150-151A CD OE cassette into *hph* marked expression vector used for transformation. |
| pBJ262 | Empty Gateway Cas9 expression vector marked with *hph* used for generating final expression constructs used for transformation. |
| pSB238 | Empty Gateway expression vector marked with *hph* used for generating final OE constructs used for transformation. |
| pDI71 | mCherry OE pDONR plasmid: pACT1:-mCherry-tCATB used as PCR template |
|  | with NA139/140 to amplify empty OE cassette used to insert *RPD3* ORF. |

**Table S3.**
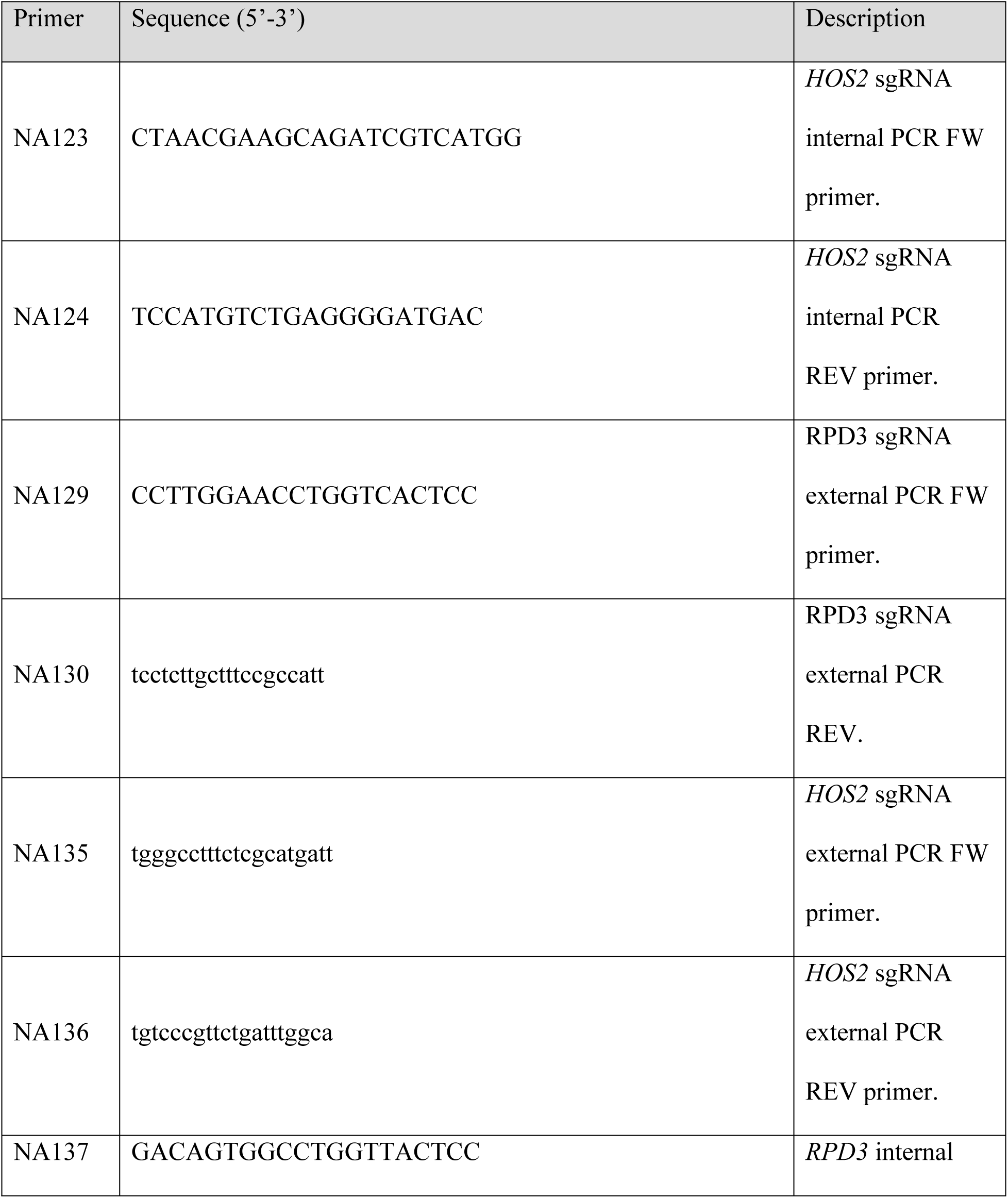
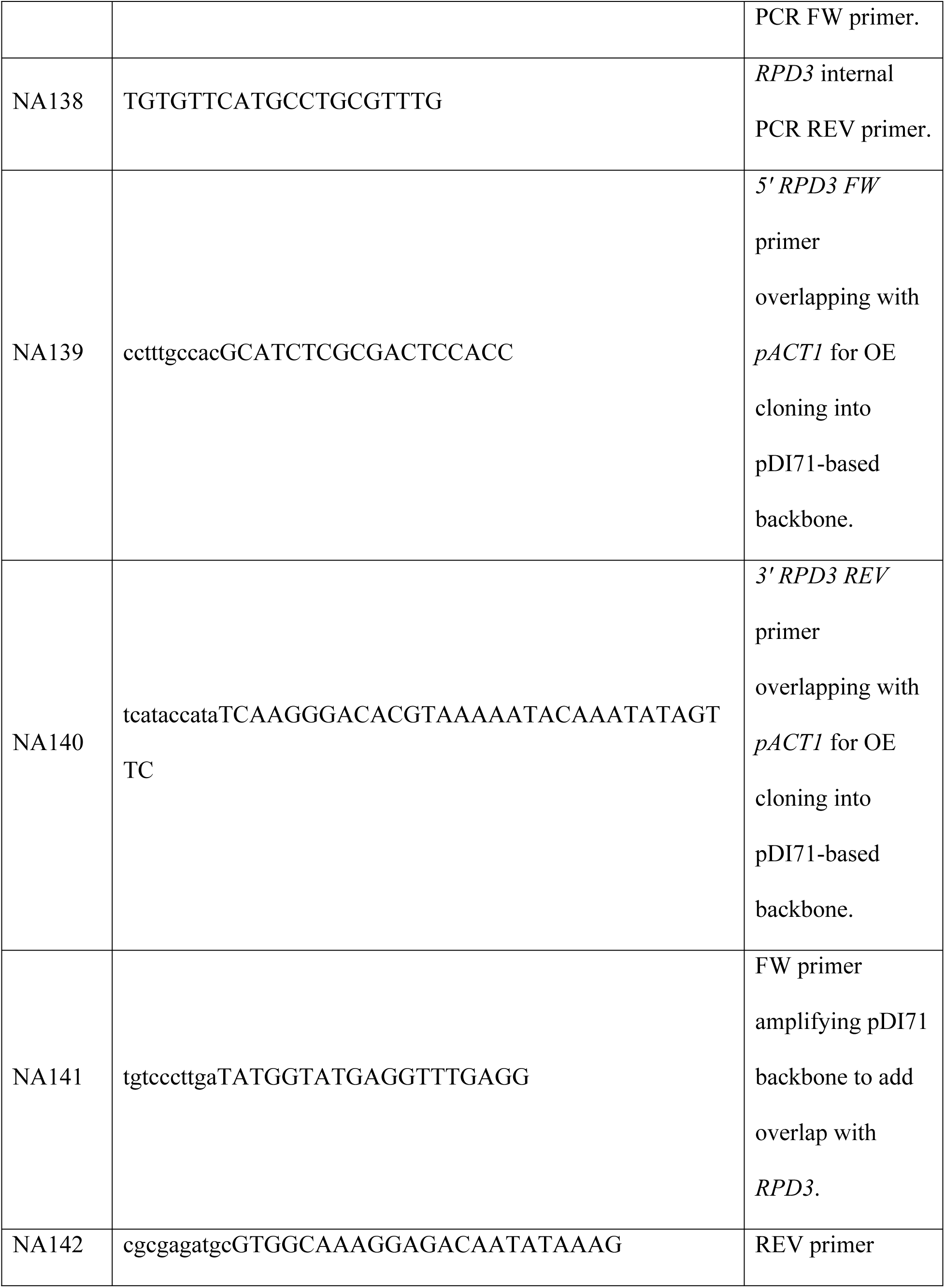
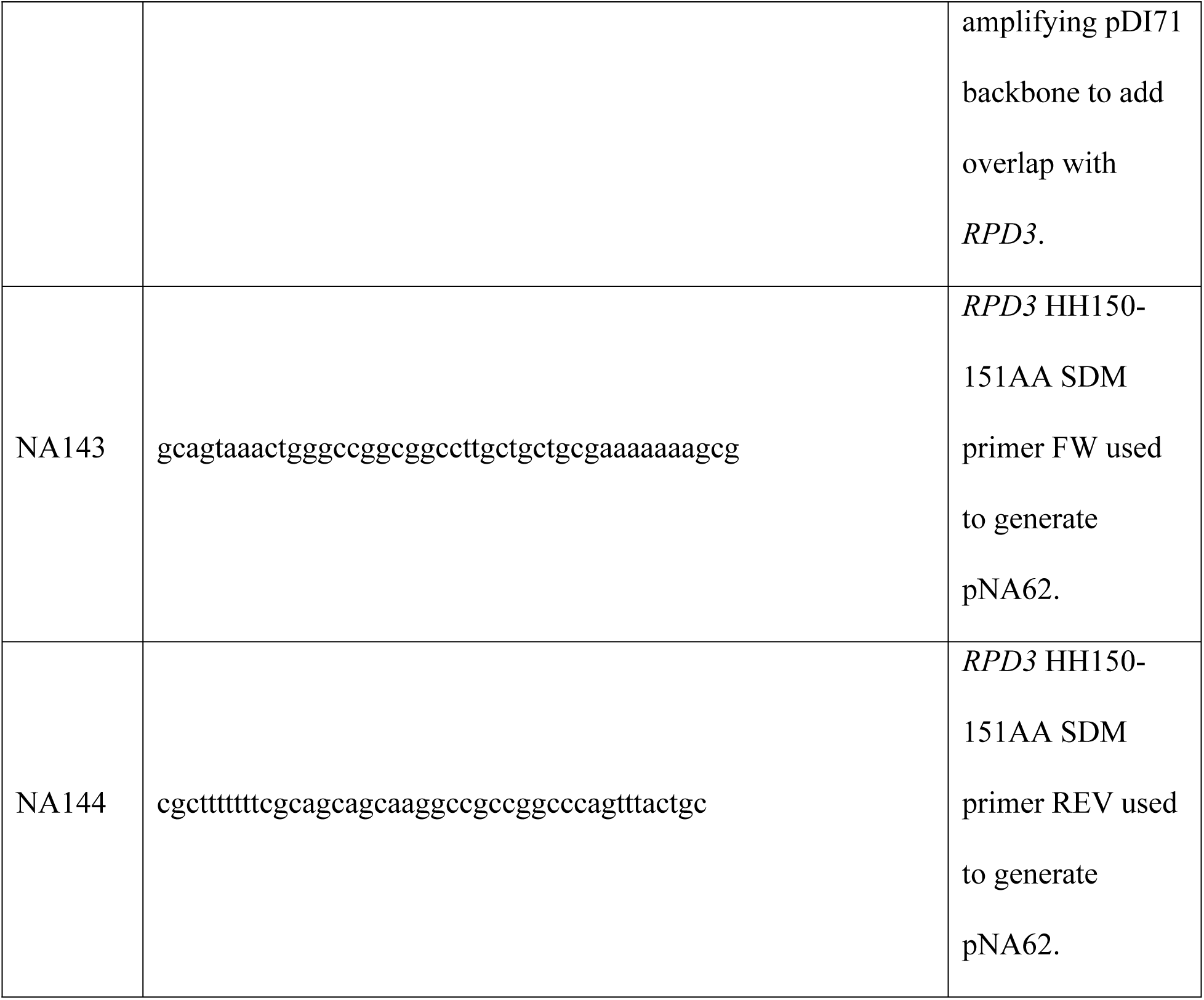
Primers used in this study.

**Table S4.**
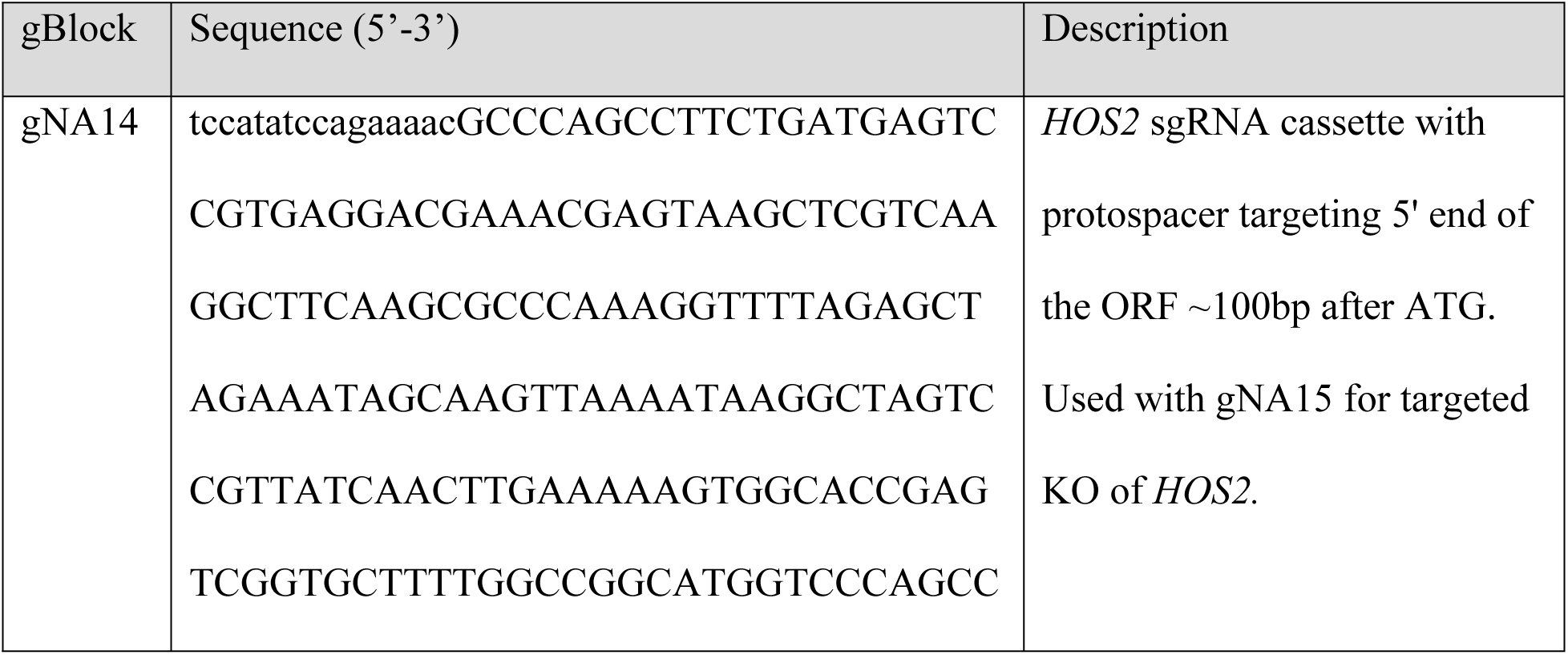
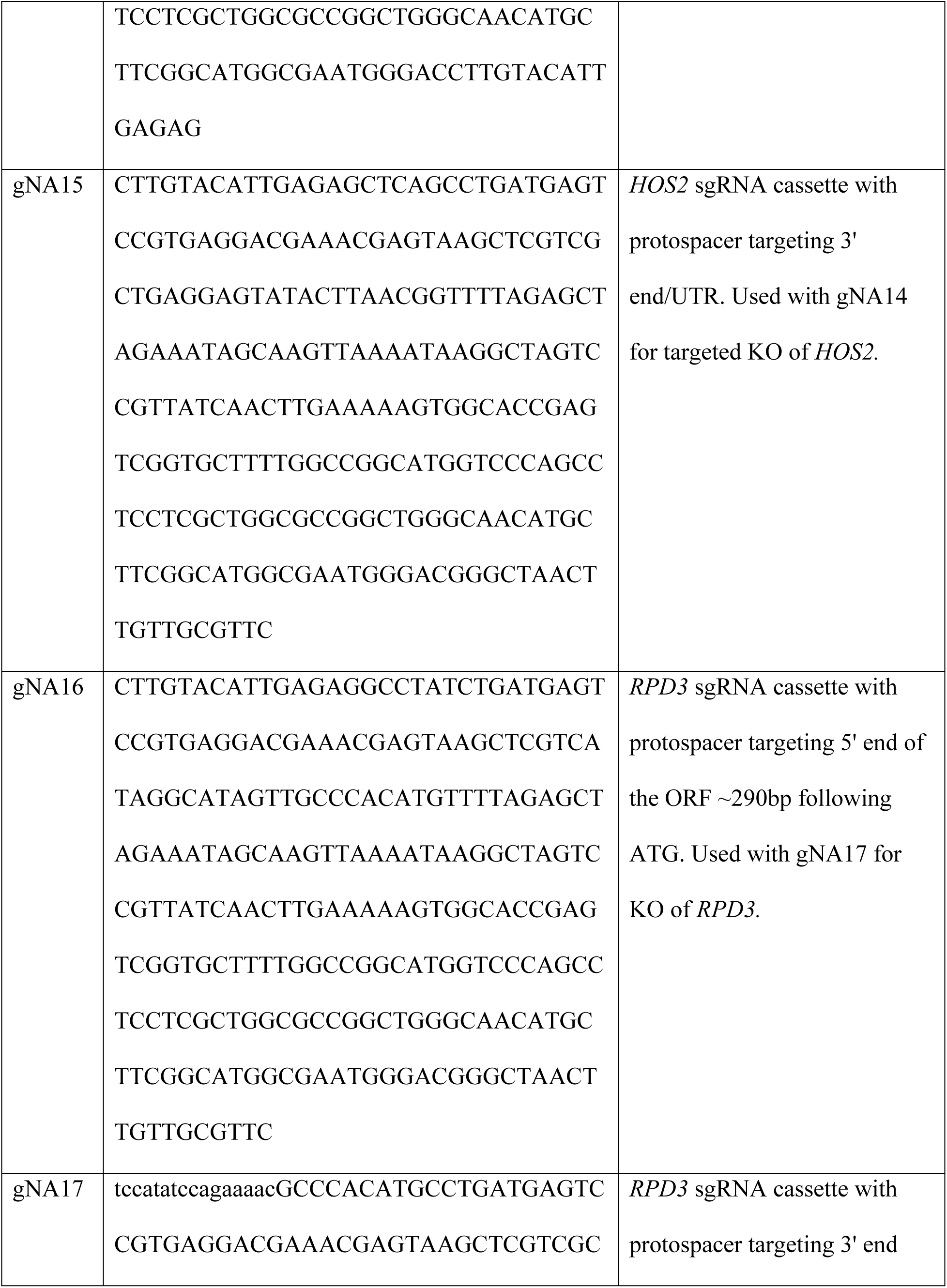
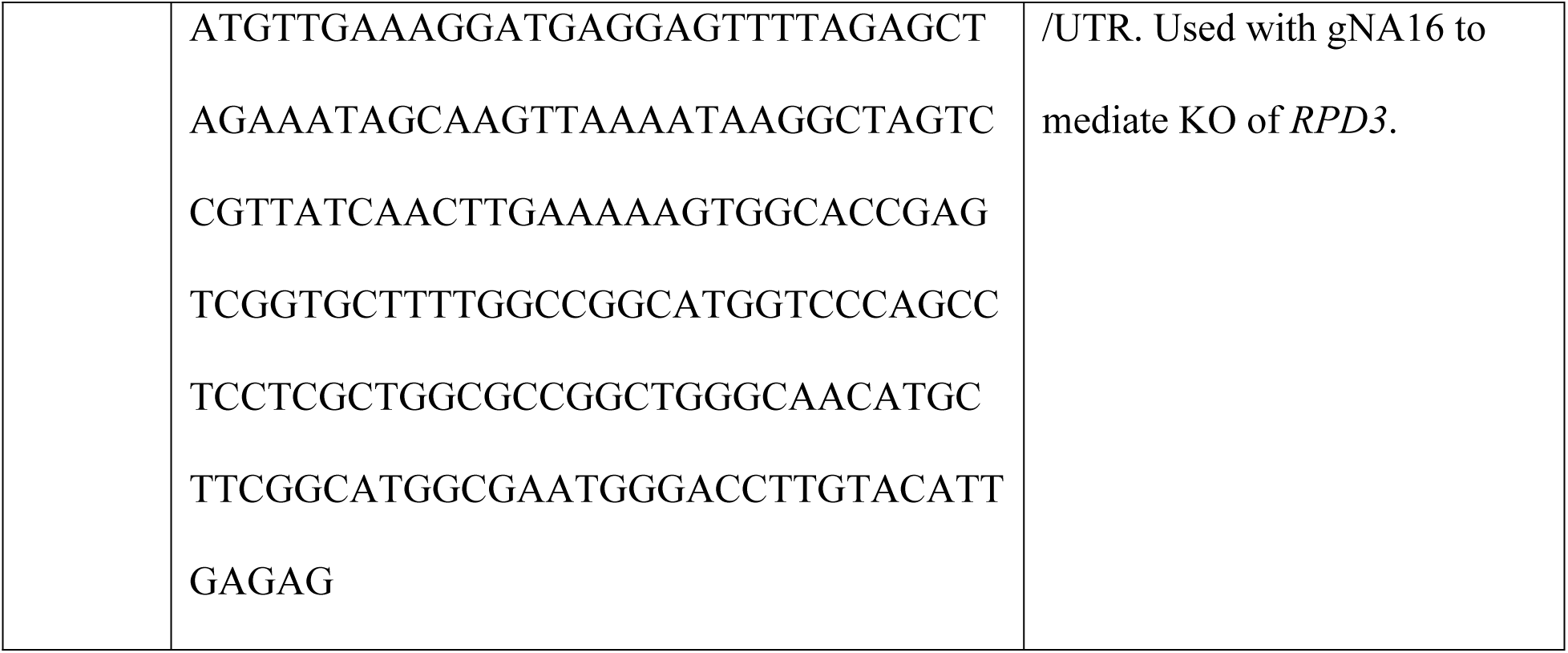
gBlocks used in this study.

**Figure S1:** Comprehensive assessment of classic HDAC phylogeny in fungi. Phylogenetic tree of classic HDAC-encoding genes in a panel of 41 fungal proteomes encompassing human fungal pathogens as well as environmental, entomopathogenic, and phytopathogenic fungi. Table S5 contains the full list of gene names and their corresponding genomes. Genes found in the G217B laboratory strain used in this study are bolded.

**Figure S2:** Entinostat induces hyphal gene expression across different conditions at 37 °C and drives filamentation through known morphology TFs. Scatterplots comparing differential RNA-seq signal in all G217B transcripts between 50 µM of Entinostat on day 1 and day 2 (A), 10 µM of Entinostat on day 1 and day 2 (B), 10 µM and 50 µM of Entinostat on day 2 (C), and 2-day RT shift and steady-state hyphae (D) (Gilmore *et al.* 2015 [18]). Venn diagrams comparing the regulons of 2-day RT (short RT), 50 µM Entinostat, and steady-state hyphae (previously published in Gilmore *et al.* 2015 [18]) to identify shared and unique upregulated (E) and downregulated genes (F).

**Figure S3:** Validation and primary characterization of HDAC KO mutants. Colony PCR products run on a 1% Agarose gel to identify successful KO isolates for *RPD3* (A) and *HOS2* (B) The top half of each gel contains PCR products amplified with a primer pair external to the sgRNA sites that is indicated in Fig. 3A. The bottom half uses a primer pair that flanks both sides of the 5’ sgRNA cut site such that negative signal is indicative of clean KO. Coverage tracks displaying sequencing signal at the chromosomal coordinates surrounding *RPD3* for control and *rpd3Δ* strains (C), and at *HOS2* for control and *hos2Δ* strains (D). E. Plate morphology phenotypes of WT, *rpd3Δ* and *hos2Δ* strains. F. OD_600_ growth curve of WT and *rpd3Δ* strains grown for 5 days at 37 °C. Points represent the average OD_600_ values of samples in triplicate. G. OD_600_ growth curve of *rpd3Δ* passaged at 37°C for 4 days in the presence of cell wall stressors. Points represent the average OD_600_ values of samples in triplicate.

**Figure S4.** Identifying putative histidine residues required for catalytic activity of Rpd3 in *Histoplasma*. A. Alignment of the Rpd3 HDAC domain across fungal species and alongside human HDAC1 (HDAC1/124-198). *Histoplasma* G217B Rpd3 is indicated by the red asterisk. The red arrows indicate the 150H and 151H residues observed to be critical for catalytic activity in orthologous species. B. Representative micrographs of transformants yielded following transformation of WT and *rpd3Δ* strains with an empty vector or a plasmid overexpressing a Catalytic Dead (CD) variant of Rpd3.

**Figure S5.** The global gene expression signature of *rpd3Δ* is better correlated with hyphae and Entinostat than *hos2Δ*. Scatterplots comparing differential RNA-seq signal in all G217B transcripts between steady-state hyphae and the following: *rpd3Δ* at 37 °C (A) and *hos2Δ* at 37 °C (B). Venn diagrams comparing the regulons identified in 2-day 50 µM Entinostat-treated cells, *rpd3Δ* at 37*°*C, and steady-state hyphae (Gilmore *et al.* 2015 [18]) to identify shared and unique sets of upregulated (C) and downregulated (D) genes. E. Global scatterplot comparing differential RNA-seq signal as in (A-B) between 37 °C signal for *hos2Δ* cells and cells grown with 50 µM Entinostat for 2 days.

**Figure S6.** Correspondence between transcriptional activation and histone acetylation in phase-specific genes. A. Scatter plot displayed in Figure 4F indexed on TF-encoding genes in the G217B genome. The green stars point out candidate hyphal TFs that are significantly induced under one of the two Rpd3-disrupting conditions. Coverage tracks displaying fold enrichment traces for H3K9Ac signal surrounding the loci of yeast-associated genes. Red traces indicate signal in control cells for both conditions in the loci surrounding *CBP1* (B) and *YPS3* (C).

**Figure S7.** Rpd3 is required for hypoacetylation of hyphal and conidial genes at 37 °C. Coverage tracks displaying traces of genomic ChIP/input signal for pulldowns on two different histone H3 acetylation marks, H3K9Ac and H3K14Ac. 2-day 37 °C cultures of WT G217B yeast treated with DMSO or 50 µM Entinostat along with G217B WT and *rpd3Δ* strains were used for chromatin isolation and subsequent pulldown. Colored traces (H3K9Ac in red and H3K14Ac in green) depict the signal in WT or DMSO control cells while the black traces represent signal with Entinostat or in *rpd3Δ* surrounding the loci of *TYR1* (A), *GAD1* (B) and *MS95* (C).

